# mPFC axons drive cognitive control enhancement during striatal stimulation

**DOI:** 10.64898/2026.04.07.717020

**Authors:** Elizabeth M Sachse, Evan M Dastin-van Rijn, Jon P Bennek, Michelle C Buccini, Megan E Mensinger, Bradley Angstadt, Francesca A Iacobucci, Manuel Esguerra, Alik S Widge

## Abstract

Deep brain stimulation (DBS) of the ventral internal capsule/ventral striatum (VCVS) can alleviate symptoms of mental illness and may work in part by improving cognitive control, a feature of healthy decision making. The human VCVS DBS target and its mid-striatum analog in our preclinical rodent model contain both cortical axons and local striatal cell bodies. However, the specific neural components that stimulation acts on to mediate cognitive effects remain unclear. We addressed this by delivering high frequency optogenetic stimulation (“opto-DBS”) to either medial prefrontal cortex (mPFC) axons or local mid-striatal (midSTR) neurons in a rodent Set-Shift task. Opto-DBS of mPFC axons reduced response times, effectively replicating the cognitive enhancement observed in a previous study that used electrical stimulation. Conversely, we observed cognitive impairment from sustained (>10 min) opto-DBS of midSTR neurons. In addition, the cognitive benefit from axonal opto-DBS exhibited time-dependent declines that were not observed with electrical stimulation. The improvement in response times declined within a session and was associated with attenuated magnitude of local midSTR evoked-response potentials. Across-testing days, we linked this decline to diminished mPFC responsiveness, evidenced by a reduction in the post-DBS functional strength of the circuit, suggesting a neuroplastic-like mechanism. These results demonstrate that mPFC-originating axons, rather than local neurons, are the primary drivers of cognitive control enhancements from electrical stimulation, providing insight into therapeutic mechanisms of VCVS DBS.

## Introduction

Deep brain stimulation (DBS) of the ventral internal capsule/ventral striatum (VCVS) can relieve symptoms in people suffering from obsessive compulsive disorder (OCD), major depressive disorder (MDD) [1,2], and addiction [3]. This promising therapy may work in part by improving cognitive control, a sub-component of flexible decision-making that is often dysfunctional in these conditions [4,5]. Prior work in OCD and MDD patients, as well as individuals with no psychiatric diagnosis, has shown that VCVS DBS improves response times (RTs) on cognitive control tasks [6,7]. Follow-on work has demonstrated that RT improvements reflect improvement in evidence processing and conflict monitoring, i.e. that they are enhancements of specific cognitive control functions [8]. However, a key unanswered question is which neural components within VCVS circuitry are primarily engaged by DBS to exert its behavioral effects. Without knowing how DBS impacts brain circuitry, physiologically informed optimization of this therapy is infeasible.

The VCVS target is composed of many axon bundles projecting from cortical areas to downstream basal ganglia regions like the thalamus (the ventral capsule or “VC” component), in addition to the local cell bodies of the ventral striatum (the “VS”) [9–11]. The DBS electric field hits both of these white and gray matter components, as well as smaller axon fascicles that travel within the striatum instead of the main capsule [12]. These cortico-striato-thalamo-cortical (CSTC) circuits are known to encode different cognitive control components (reward, attention, action-selection, [13–15]) and show impaired function in several psychiatric illnesses [16–19], therefore any one of these regions could be key to the effect of DBS.

In prior work, the magnitude of the RT improvement from DBS depended on the specific stimulation location, with more dorsal electrode contacts being more effective than ventral ones [7]. Stimulation at dorsal sites might engage more “VC” e.g. white matter axons than “VS”, or they could be closer to relevant striatal circuitry, for example the caudate nucleus which is associated with cognitive and behavioral flexibility [20–22]. VCVS DBS-driven increases in theta oscillations in PFC regions [7,23] and other past work [24–26] suggest axons may be the key components. These mechanistic questions are difficult to answer in humans given limited neuron/circuit-specific tools and testing constraints. Animal models of DBS can overcome these barriers. Past work developed a rodent model of VCVS DBS leveraging human-rodent homology of VCVS circuitry and the cognitive control behavioral effect. As in humans, DBS-like stimulation of the rat mid-striatum (midSTR) reduces RTs on a cognitive control task [8]. This effect is robust, can be elicited with unilateral stimulation, and has similar effect sizes for constant vs. intermittent stimulation [27,28]. Like the VCVS, the midSTR target contains axons projecting from PFC regions as well as local striatal neurons, and both could be responsible for stimulation’s cognitive control effect. Although the topology of PFC projections to striatum and other downstream structures are analogous in humans and rats [29], rats do not have a true internal capsule [10,30]. Instead, mPFC axons projecting to and through the midSTR are intermixed with local cell bodies, making it even more challenging to target electrical stimulation to one of these neuronal components. Optogenetics enables precise circuit/cell-specific targeting of stimulation and can be used to dissect electrical stimulation paradigms [31–34] .

Here, we used optogenetics to target high-frequency stimulation, “opto-DBS”, to either mPFC axons or local striatal neurons and assess effects on cognitive control. Opto-DBS of mPFC axons, but not midSTR neurons, improved cognitive control (reduced RTs), replicating the effects of electrical stimulation. Unlike electrical stimulation, opto-DBS of both targets appeared to have time-dependent changes in the behavioral effect. Within a day, i.e. over the course of one testing session, mPFC-axon opto-DBS led to a reduction in the RT improvement as well as evoked neural activity in the midSTR. Across testing days, the decline in axonal opto-DBS’s effects was linked to a reduction in mPFC activity. These temporal changes may be due to changes in opsin function, neuroplasticity mechanisms, or inherent differences in electrical vs. photonic stimulation. Overall, these results suggest that the improvement in cognitive control from midSTR electrical stimulation is driven primarily by mPFC-originating axons versus local neurons. This is an important step towards understanding the mechanisms by which VCVS DBS improves cognitive control, an essential part of refining existing psychiatric DBS therapies.

## Methods

### Animals

38 adult Long Evans rats (20 male, 18 female, see below for power analysis) were obtained from Charles River Laboratories (Wilmington, MA), and pair-housed on a 14:10 light/dark cycle. Rat pairs were randomly assigned to one of four groups: mPFC-Chronos, mPFC-control, midSTR-Chronos, or midSTR-control. After acclimation, rats were handled for 5 min/day for at least 5 days to familiarize them with experimenters, prior to undergoing surgery. Rats were given ad libitum access to standard food and were at least three months old (adulthood) prior to undergoing surgery. Post-surgical recovery, to motivate task performance, rats were food restricted to maintain 85 to 90% of their starting body weight. Supplemental food was provided if task rewards were insufficient to meet their caloric needs, or if dominance hierarchies between pair-housed rats led to unequal food consumption. All experiments were approved by the University of Minnesota Institutional Animal Care and Use Committee (protocol 2402-41825A or 2104-39021A).

### Set-Shift Task

We used the Set-Shift task to measure cognitive control (Figure 1A), as in previous studies [8,27,35,36]. The task was run in operant chambers (Lafayette Instruments) containing three illuminable nose-pokes with infrared detectors, a food dispenser on the opposite wall to deliver sucrose pellets (45mg, BioServe) enclosed in sound-proof boxes (Med Associates). Task stimuli were controlled by *pybehave*/OSCAR software [37,38]. Rats began Set-Shift training after recovery from surgery. Rats were trained to adjust their responses between a cue-driven “Light” rule and a spatial “Side” rule. To initiate a trial, rats had to poke the illuminated middle nose-poke, then poke one of the two peripheral nose-pokes, one of which was illuminated. Rats were reinforced with a single reward pellet for each correct response. After rats made five consecutive correct choices, the rule switched to the other dimension without any cue, and rats had to shift their behavior to continue receiving rewards. The task contained eight “blocks” defined as the sequential trials on a single rule regardless of accuracy, requiring rats to shift rules seven times per test session.

**Figure 1:**
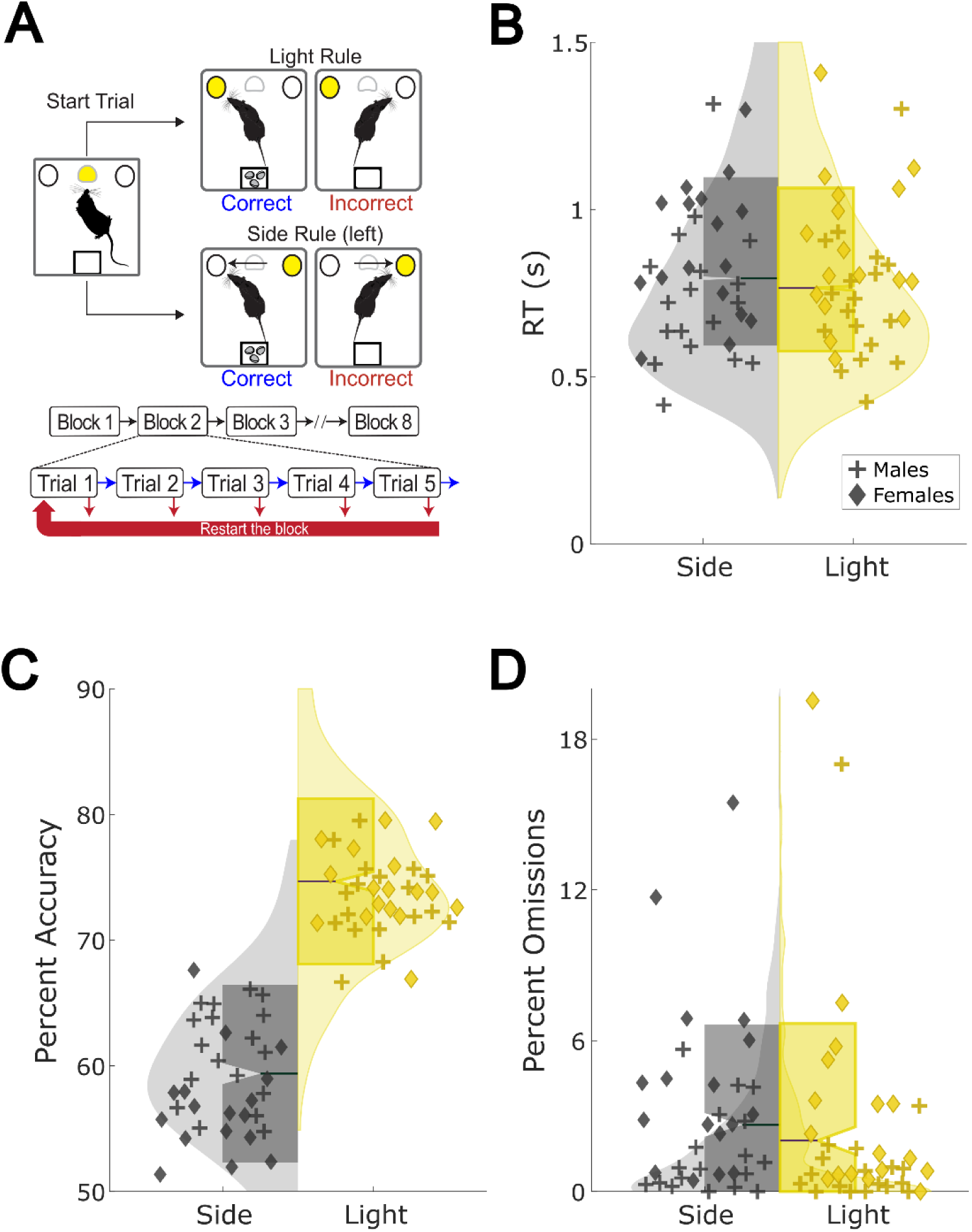
The rodent Set-Shift task as a translational model of cognitive control. **A)** The Set-Shift task. Rats initiate a trial by poking the middle nosepoke and then must nosepoke either an illuminated hole (Light rule) or ignore the light and always nosepoke on a specific side of the chamber (Side rule). The rules are not cued and shift to a new rule after the rat has sequentially completed 5 correct trials of the current rule. Errors reset at the beginning of the current 5-trial block. **B)** DBS-OFF response times (RT) for all rats (N = 35) on Side (gray) and Light (yellow) rule trials. Scatter points represent median OFF RT for individual rats (triangles = female, crosses = males). Distributions illustrated by violins were computed over all trials. **C)** OFF session distributions of percent accuracy for all rats computed over all sessions. Scatters represent mean OFF accuracy for each rat. **D)** OFF session distributions of percent omissions for all rats computed over all sessions. Scatters represent mean OFF accuracy for each rat.

To move on from training to optic cable habituation, rats completed 8 to 12 training sessions of habituation to a branched fiber-optic stimulation cable (custom-designed by Doric Lenses and connected Doric Lenses fiber-optic rotary joint) for a minimum of 4 training sessions before beginning testing. We used four test session protocols with different rule orders to prevent predictability of the next trial. Each protocol started and ended with a sequence of 10 trials where rewards were given randomly, independent of the rats’ choices, to ensure the rat had no a priori rule information before the start of the first block. Rule sequences within protocols were counterbalanced so that all possible rules were presented with equal frequency. Protocol order was balanced with stimulation conditions, and was the same for each rat, except when a session could not be run (due to equipment malfunction or abnormal testing conditions). Protocols missed due to these conditions were appended to the end of the remaining sessions.

### Surgery for Chronos transfection and implantation of bilateral optic fibers

Rats were given preoperative analgesia, anesthetized with isoflurane, and mounted on a stereotaxic frame. Tissue was incised and resected over the target areas. Craniotomies were drilled at virus injection sites and fiber implantation sites and for anchor screws. Rats were injected with an optogenetic virus in either mPFC or midSTR.Because we wanted to reproduce electrical DBS parameters as closely as possible, we chose to use the Chronos opsin as it has been shown to follow high-frequency stimulation [39–42]. mPFC rats were injected with 0.5 µL of either AAV5-CaMKII-Chronos-GFP or AAV5-CaMKII-GFP bilaterally in three mPFC subregions (+2.8 AP, +/-0.6 ML, -5, -3.7, and -2.3 DV) to target glutamatergic projection neurons across a large dorsal-ventral span of bilateral mPFC. midSTR rats received 0.25 µL of either AAV5-hSyn-Chronos-GFP or AAV5-hSyn-GFP bilaterally in two midSTR subregions (+1.4 AP, +/- 2.0 ML, -5.7 and -6.1 DV) to target all neuronal cell bodies within the midSTR. All rats were implanted bilaterally with 400 µm, 0.5 NA, 7mm long FC optic fibers (RWD Life Science) to deliver optical stimuli to the same midSTR area used in prior electrical DBS studies [8]. Fibers were affixed to the skull and screws with metabond and dental cement. After surgery, rats were returned to their home cages and recovered for 1 to 2 weeks before beginning food restriction for training on the Set-Shift task. Viral expression was allowed to develop for at least 5 weeks during surgical recovery and the Set-Shift training period before the delivery of any optical stimulation. Chronos activation with 470 nm light stimulation was confirmed in a separate cohort of rats that were used for the electrophysiology experiments (see below).

### DBS-like optical stimulation during set-shift

After sufficient cable habituation, rats began Set-Shift testing sessions. Optical stimulation (“Opto-DBS”) was delivered using a PC-controlled StimJim stimulator that controlled a LED driver connected to a 470 nm LED (ThorLabs). For “ON” sessions, Opto-DBS (130 Hz, 2 ms pulse width 4V LED driver input) was delivered to bilateral mid-striatum through the duration of the task. 130 Hz is the same frequency used for electrical stimulation in prior work [8] and common in clinical DBS [43]. For sham (“OFF”) sessions, rats were connected to cables without light delivery. Each rat underwent 16 sessions total (8 ON, 8 OFF), in a pseudorandom, counterbalanced order.

### Electrophysiology experiments

Electrophysiology recordings were used to verify Chronos-driven activation of mPFC or midSTR neurons and to determine if changes in the behavioral effects of mPFC-axon opto-DBS over time are accompanied by changes in neural activity in these regions. All animal and surgical procedures were identical to the methods described above for Set-Shift experiments, unless specifically noted below.

#### mPFC-axon opto-DBS experiment

4 adult Long Evans rats (2M, 2F), were injected with AAV5-CaMKII-Chronos-GFP (N = 4, same targets and volumes as above) and implanted with in-house built opto-microwire electrodes (3 to 4 wires per site, 50 or 75 µm platinum-iridium, AM Systems) in bilateral mPFC and midSTR, connected to a 32-channel connector (Omnetics) to enable local field potential (LFP) recordings. MidSTR electrodes were adjacent to optic fibers for stimulation. Surgical procedures were the same as above, except that viral transfection and electrode implantation occurred in separate procedures to preserve recording quality. Ground and reference wires were tied to cerebellar skull screws. Rats were single-housed post-electrode surgeries to prevent chewing of connectors but were socialized with experimenters or their former cage mate a minimum of twice per week.

After surgical recovery, rats were food-restricted to ∼95% of their starting body weight to mimic Set-Shift restriction conditions. Rats were tested weekly for the presence of optically evoked response potentials (ERPs) in midSTR and mPFC, indicating successful viral transfection and electrode targeting. Main recording sessions began once ERPs in all four recording sites were present or if rats were >= 60 days post-transfection and there was no expected improvement in ERP detection. All 4 rats tested had at least one striatal site with an ERP. Rats underwent 8 recording sessions, each comprised of 1) a pre-DBS bilateral sweep over amplitudes for ERPs (2s 5 Hz pulse trains, 4s OFF interval, 50 trains per amplitude of 0 to 5V LED driver inputs, in steps of 1V), 2) 30min of 130 Hz task-like bilateral opto-DBS, and 3) a post-DBS bilateral ERP amplitude sweep. Throughout the session, rats were in an open field arena (31 x 26 x 29 cm) and tethered to the same type of optic-fiber stimulation cable used for Set-Shift opto-DBS and a 32-channel recording headstage+ cable (Intan Technologies, Los Angeles, CA), that were both connected to an assisted fiber-optic & electrical rotary joint (Doric Lenses). Sessions were run on the same schedule as Set-Shift opto-DBS sessions (ON/recording session, OFF/no session, ON/recording…). Session events and optical stimuli were controlled using *pybehave*/OSCAR software and a StimJim-controlled LED driver, like Set-Shift experiments described above. LFPs were recorded (1.4 Hz high-pass, 7600 kHz low-pass (open) sampled at 30 kHz), with an OpenEphys acquisition system (Atlanta, Georgia) in conjunction with additional digital acquisition systems (National Instruments, Austin, Texas) to communicate between *pybehave*/OSCAR, the StimJim/LED driver, and OpenEphys systems.

### Histology

After completion of Set-Shift sessions or electrophysiology experiments, rats were deeply anesthetized with isoflurane, euthanized (Euthasol, 100 mg/kg), and transcardially perfused with PBS followed by 4% paraformaldehyde. Brains were then extracted, post-fixed in PFA, cryoprotected in sucrose, and sectioned at 30 µm on a cryostat. Sections were mounted on slides, Hoechst or DAPI-labeled, and imaged on a fluorescent microscope (Keyence) to verify viral transfection and fiber/electrode placements. Successful viral transfection and targeting was based on raw GFP expression in mPFC and/or midSTR, depending on the experimental group. Fibers were considered on-target if the fiber tips resided in the midSTR region previously defined in electrical stimulation experiments [8]. Because here we used optical stimulation, we targeted fiber tips to ∼300 µm dorsal to the electrical target, to illuminate the same area at the electrode tip. Fiber placements were visualized using the ScatterBrain toolbox [44], which models the area illuminated based on optical parameters such as numerical aperture and fiber diameter (Figures 2B and 3B).

**Figure 2:**
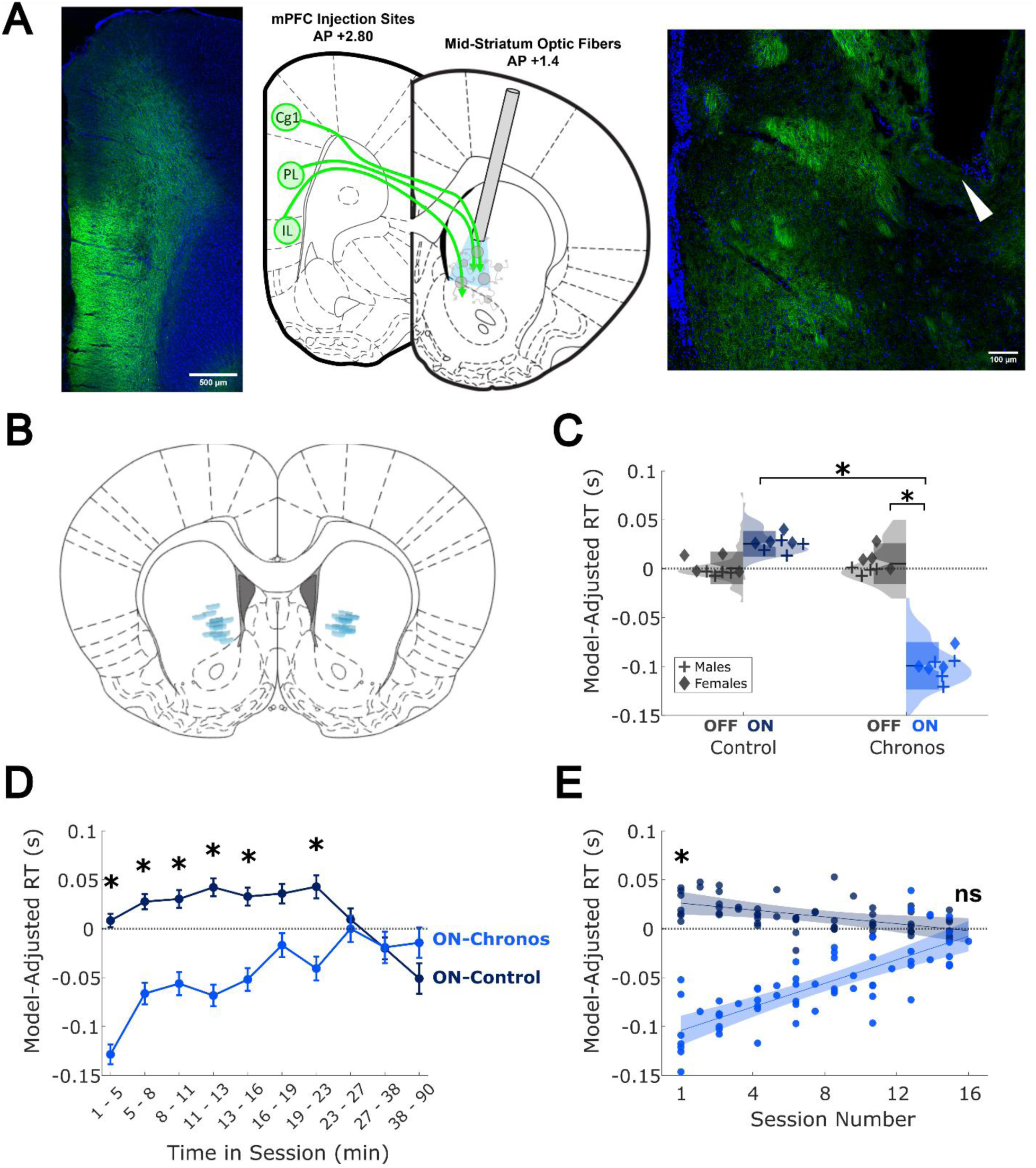
Opto-DBS of mPFC axons improves cognitive control. **A)** AAV5-CaMKII-Chronos-GFP or control was injected in the mPFC and optic fibers were implanted in the midSTR to enable opto-DBS of mPFC axons that project to and through the midSTR. Fluorescent images show GFP+ mPFC cell bodies (left) and axons (right) at coordinates corresponding to atlas slices in the middle. White triangle indicates fiber tip in midSTR. **B)** Fiber placement in midSTR, illustrating the area illuminated by the fiber tip. Atlas slice corresponds to mean AP coordinate across mPFC rats. **C)** Distributions of model-adjusted RT for control (left) and Chronos (right) rats, computed over trials. Scatter points represent individual rat median adjusted-RTs (triangles = females, pluses = males). Axonal opto-DBS significantly reduced RTs compared to both the same rats with opto-DBS OFF (right *, p = 0.00027) and to opto-DBS ON in rats with a control virus (p = 0.00015). See the Methods statistical analysis section for a description of units of y-axis. **D)** Effect of opto-DBS on adjusted RT during a session. Scatter points represent mean RT change at that time bin in the session, for ON-Chronos (light blue) and ON-control (dark blue) groups ± SEM. The RT reduction from opto-DBS declines throughout a session, so that ON-Chronos and ON-control RTs are not significantly different by bin 8 (* indicates p < 0.05, see Table 1 in Supplement). **E)** Effect of opto-DBS on model-adjusted RT in Chronos (light blue) vs. control (dark blue) rats across sessions. Scatter points represent individual rat’s mean RT change at that session. Shaded regions shows 95% confidence intervals. At session 1, ON-Chronos RTs are significantly reduced by DBS compared to ON-controls (p = 0.00014). But by session 16, there is no significant difference (p = 0.7182).

**Figure 3:**
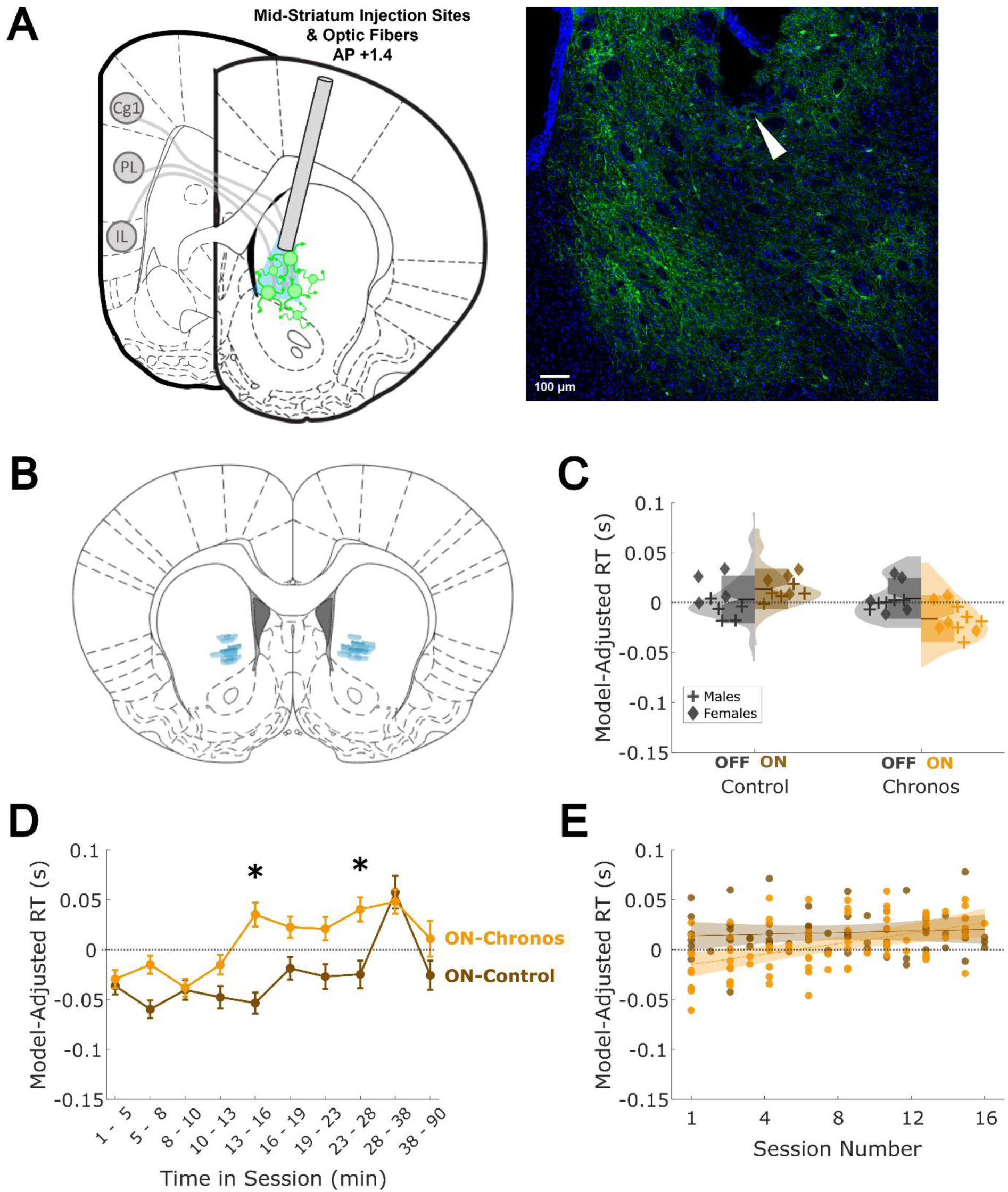
Sustained (> 10 minutes) opto-DBS of midSTR impairs cognitive control. **A)** AAV5-hSyn-Chronos-GFP or control was injected in the midSTR and optic fibers were implanted at the same coordinates to enable opto-DBS of local striatal neurons. Image of GFP+ fluorescence in midSTR neurons corresponds to atlas slice on the left. White triangle points to fiber tip. **B)** Fiber placement in midSTR, illustrating the area illuminated by the fiber tip. Atlas slice corresponds to mean AP coordinate across rats. **C)** Distributions of model-adjusted RT for control (left) and Chronos (right) rats. Scatter points represent individual rat median adjusted-RTs (triangles = females, pluses = males). midSTR neuron opto-DBS did not significantly change RT compared to ON-controls (p = 0.4125). **D)** Effect of opto-DBS on model-adjusted RT throughout a session. Scatter points represent mean RT change at that time bin in the session, for ON-Chronos (light orange) and ON-control (dark orange) groups ± SEM. midSTR neuron DBS began to slow RTs after ∼10-15 minutes of DBS during Set-Shift (* indicates p < 0.05, see Table 2 in Supplement). **E)** Effect of opto-DBS on model-adjusted RT in Chronos (light orange) vs. control (dark orange) rats across sessions. Scatter points represent individual rat’s mean RT change at that session. Shaded regions shows 95% confidence intervals. There were no differences between ON-Chronos and ON-control at session 1 (p = 0.4125), nor at session 16 (p = 0.8000).

**Table 1:**
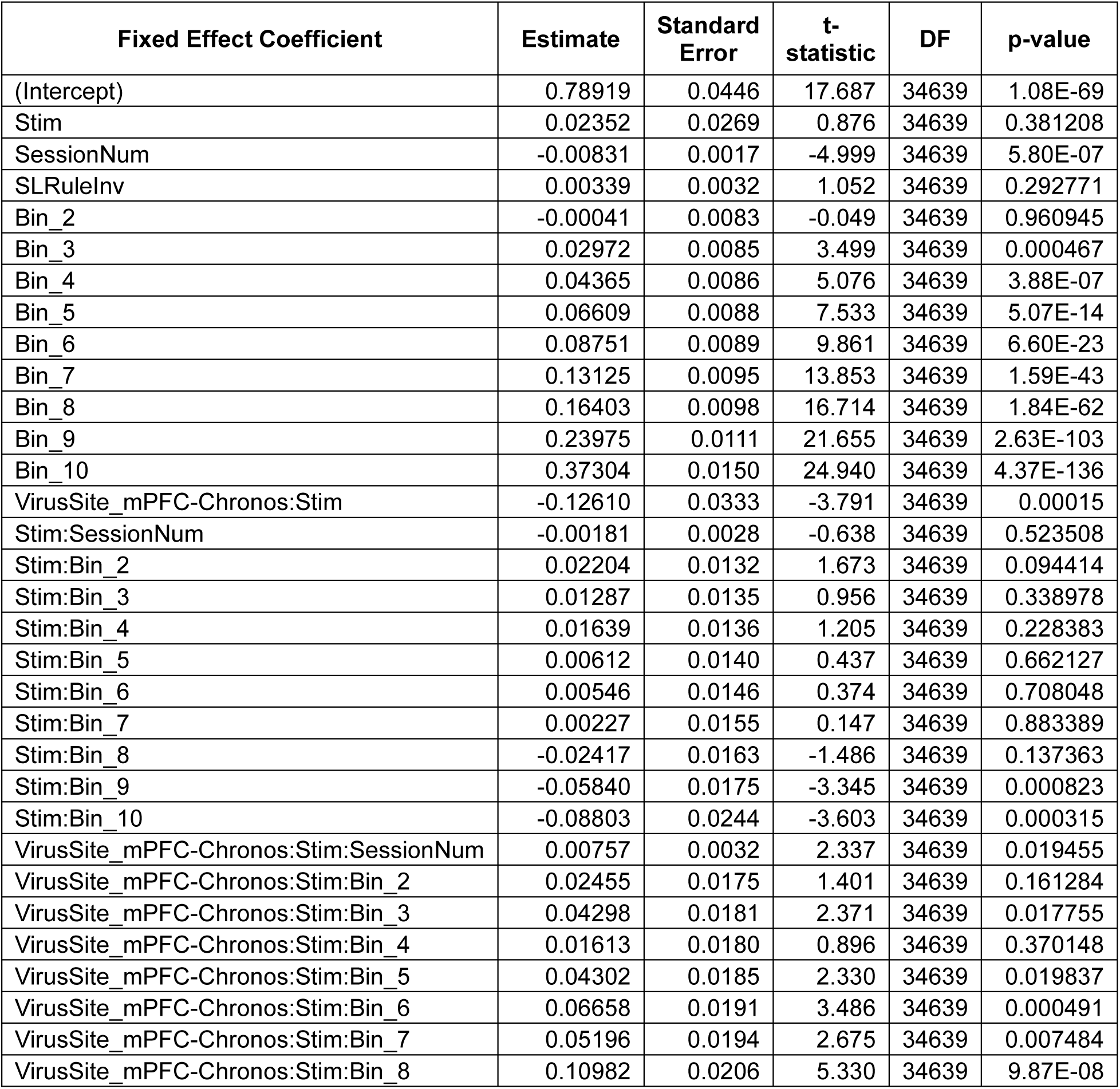
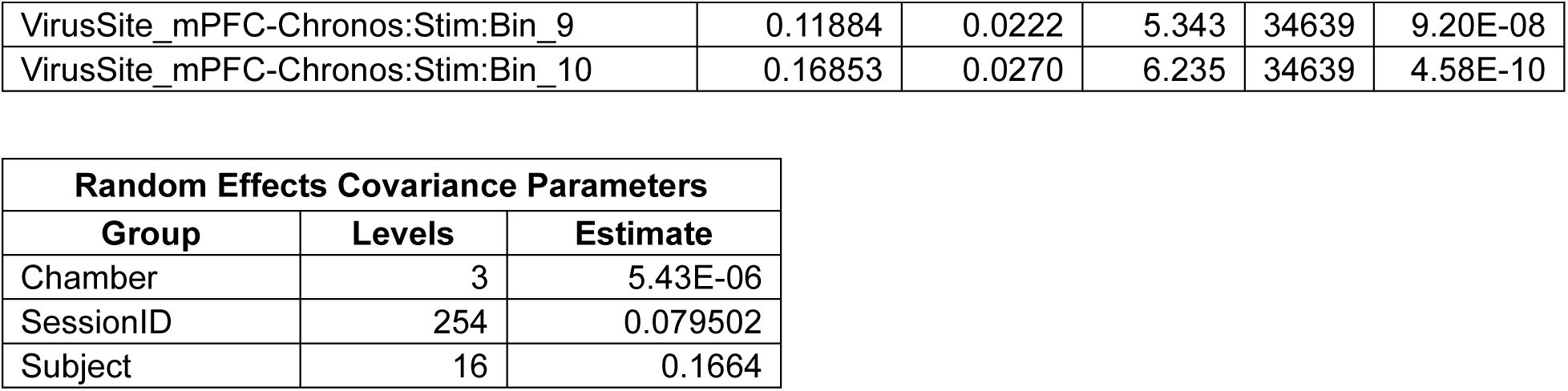
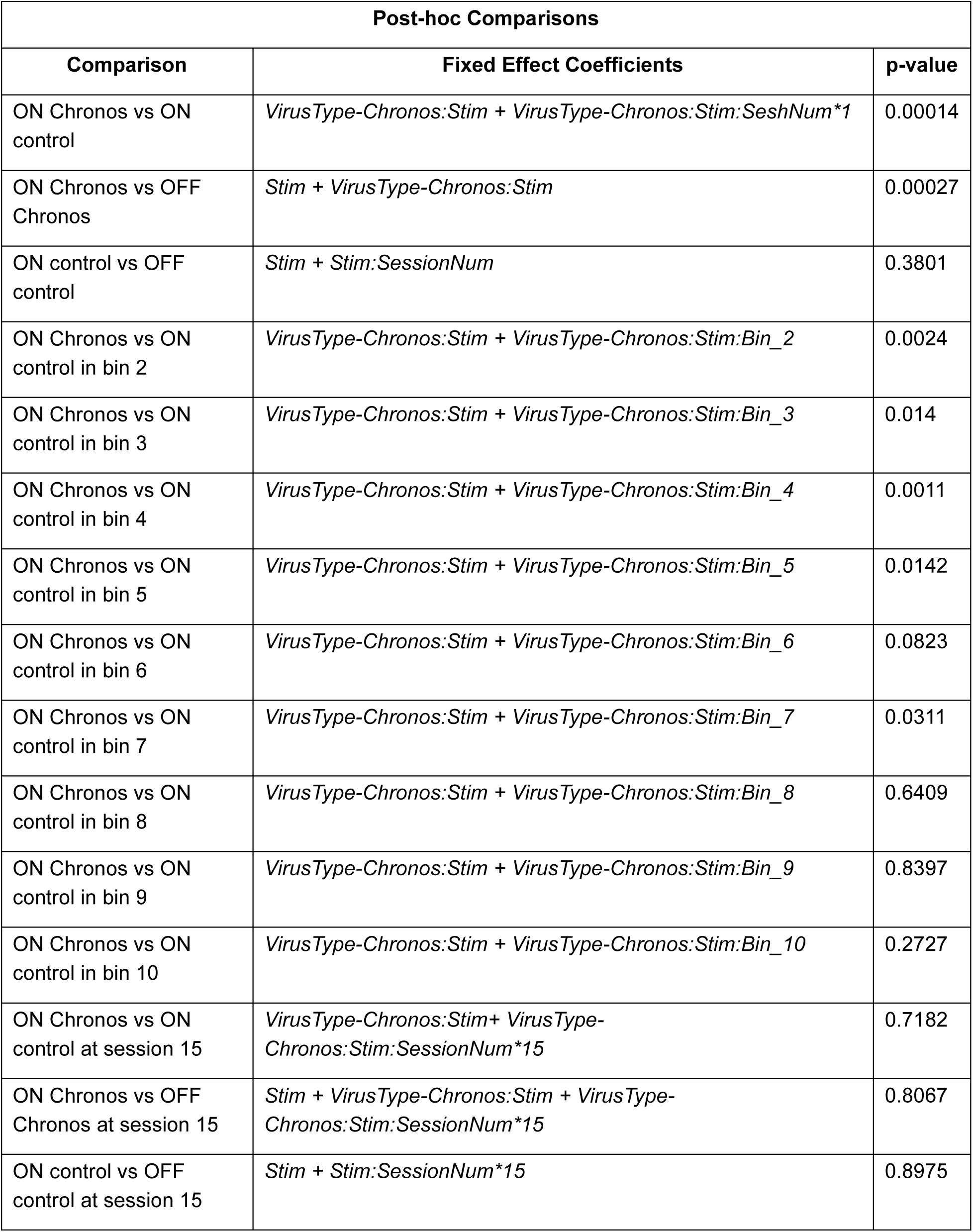
Generalized-linear-model coefficients for RT in mPFC group. Regression coefficients for response time (RT) in seconds in the mPFC-axon opto-DBS experiment from a generalized linear mixed-effects model (GLM), with a gamma distribution and identity link function. **Formula**: *RT ∼ 1+ Stim + BinnedTIS + SessionNum + RuleType + Stim:BinnedTIS + Stim:SessionNum + Stim:VirusSite + Stim:VirusSite:SessionNum + Stim:VirusSite:BinnedTIS + (1|Chamber) + (1|SessionID) + (1|Subject)*

**Table 2:** Generalized-linear-model coefficients for RT in midSTR group. Regression coefficients for RT in seconds in the midSTR-neuron opto-DBS experiment from a GLM with a gamma distribution and identity link function. **Formula**: *RT∼1+ Stim + BinnedTIS + SessionNum + RuleType + Stim:BinnedTIS + Stim:SessionNum + Stim:VirusSite + Stim:VirusSite:SessionNum + Stim:VirusSite:BinnedTIS + (1|Chamber) + (1|SessionID) + (1|Subject)*

| Fixed Effect Coefficient | Estimate | Standard Error | t-statistic | DF | p-value |
| --- | --- | --- | --- | --- | --- |
| '(Intercept)' | 0.81274 | 0.0474 | 17.139 | 41232 | 1.29E-65 |
| 'Stim' | 0.01093 | 0.0309 | 0.354 | 41232 | 0.72356505 |
| 'SessionNum' | -0.00536 | 0.0018 | -2.964 | 41232 | 0.00304001 |
| 'SLRuleInv' | 0.00256 | 0.0032 | 0.790 | 41232 | 0.42952687 |
| 'Bin_2' | 0.03523 | 0.0081 | 4.371 | 41232 | 1.24E-05 |
| 'Bin_3' | 0.07922 | 0.0083 | 9.490 | 41232 | 2.44E-21 |
| 'Bin_4' | 0.11856 | 0.0086 | 13.731 | 41232 | 8.27E-43 |
| 'Bin_5' | 0.14211 | 0.0087 | 16.247 | 41232 | 3.58E-59 |
| 'Bin_6' | 0.16748 | 0.0090 | 18.712 | 41232 | 8.33E-78 |
| 'Bin_7' | 0.21377 | 0.0094 | 22.782 | 41232 | 3.50E-114 |
| 'Bin_8' | 0.24983 | 0.0098 | 25.466 | 41232 | 5.96E-142 |
| 'Bin_9' | 0.27201 | 0.0102 | 26.548 | 41232 | 5.35E-154 |
| 'Bin_10' | 0.38773 | 0.0124 | 31.357 | 41232 | 2.51E-213 |
| 'VirusSite_midSTR-Chronos:Stim' | -0.03025 | 0.0369 | -0.820 | 41232 | 0.412471494 |
| 'Stim:SessionNum' | 0.00029 | 0.0032 | 0.089 | 41232 | 0.92882147 |
| 'Stim:Bin_2' | -0.03248 | 0.0136 | -2.387 | 41232 | 0.01698622 |
| 'Stim:Bin_3' | -0.03765 | 0.0142 | -2.659 | 41232 | 0.00783724 |
| 'Stim:Bin_4' | -0.04959 | 0.0145 | -3.421 | 41232 | 0.00062463 |
| 'Stim:Bin_5' | -0.07001 | 0.0146 | -4.800 | 41232 | 1.60E-06 |
| 'Stim:Bin_6' | -0.05034 | 0.0150 | -3.360 | 41232 | 0.00078147 |
| 'Stim:Bin_7' | -0.06488 | 0.0157 | -4.143 | 41232 | 3.43E-05 |
| 'Stim:Bin_8' | -0.07535 | 0.0168 | -4.482 | 41232 | 7.42E-06 |
| 'Stim:Bin_9' | 0.01261 | 0.0205 | 0.615 | 41232 | 0.53843793 |
| 'Stim:Bin_10' | -0.06428 | 0.0220 | -2.928 | 41232 | 0.00341852 |
| 'VirusSite_midSTR-Chronos:Stim:SessionNum' | 0.00267 | 0.0037 | 0.728 | 41232 | 0.46686390 |
| 'VirusSite_midSTR-Chronos:Stim:Bin_2' | 0.04554 | 0.0166 | 2.748 | 41232 | 0.00599763 |
| 'VirusSite_midSTR-Chronos:Stim:Bin_3' | 0.02536 | 0.0169 | 1.500 | 41232 | 0.13370211 |
| 'VirusSite_midSTR-Chronos:Stim:Bin_4' | 0.05237 | 0.0174 | 3.014 | 41232 | 0.00257563 |
| 'VirusSite_midSTR-Chronos:Stim:Bin_5' | 0.11269 | 0.0178 | 6.321 | 41232 | 2.62E-10 |
| 'VirusSite_midSTR-Chronos:Stim:Bin_6' | 0.08420 | 0.0182 | 4.632 | 41232 | 3.63E-06 |
| 'VirusSite_midSTR-Chronos:Stim:Bin_7' | 0.09528 | 0.0189 | 5.039 | 41232 | 4.71E-07 |
| 'VirusSite_midSTR-Chronos:Stim:Bin_8' | 0.12299 | 0.0202 | 6.091 | 41232 | 1.13E-09 |
| 'VirusSite_midSTR-Chronos:Stim:Bin_9' | 0.04764 | 0.0246 | 1.934 | 41232 | 0.05305957 |
| 'VirusSite_midSTR-Chronos:Stim:Bin_10' | 0.08378 | 0.0294 | 2.854 | 41232 | 0.00432059 |

| Random Effects Covariance Parameters |  |  |
| --- | --- | --- |
| Group | Levels | Estimate |
| Chamber | 4 | 7.31E-06 |
| SessionID | 304 | 0.097757 |
| Subject | 19 | 0.19222 |

| Post-hoc Comparisons |  |  |
| --- | --- | --- |
| Comparison | Fixed Effect Coefficients | p-value |
| ON Chronos vs ON control | *VirusType-Chronos:Stim + VirusType-Chronos:Stim:SeshNum\*1* | 0.4224 |
| ON Chronos vs OFF Chronos | *Stim + VirusType-Chronos:Stim* | 0.514 |
| ON control vs OFF control | *Stim + Stim:SessionNum* | 0.6934 |
| ON Chronos vs ON control in bin 2 | *VirusType-Chronos:Stim + VirusType-Chronos:Stim:Bin_2* | 0.6795 |
| ON Chronos vs ON control in bin 3 | *VirusType-Chronos:Stim + VirusType-Chronos:Stim:Bin_3* | 0.8951 |
| ON Chronos vs ON control in bin 4 | *VirusType-Chronos:Stim + VirusType-Chronos:Stim:Bin_4* | 0.5533 |
| ON Chronos vs ON control in bin 5 | *VirusType-Chronos:Stim + VirusType-Chronos:Stim:Bin_5* | 0.028 |
| ON Chronos vs ON control in bin 6 | *VirusType-Chronos:Stim + VirusType-Chronos:Stim:Bin_6* | 0.1522 |
| ON Chronos vs ON control in bin 7 | *VirusType-Chronos:Stim + VirusType-Chronos:Stim:Bin_7* | 0.0862 |
| ON Chronos vs ON control in bin 8 | *VirusType-Chronos:Stim + VirusType-Chronos:Stim:Bin_8* | 0.0158 |
| ON Chronos vs ON control in bin 9 | *VirusType-Chronos:Stim + VirusType-Chronos:Stim:Bin_9* | 0.6685 |
| ON Chronos vs ON control in bin 10 | *VirusType-Chronos:Stim + VirusType-Chronos:Stim:Bin_10* | 0.2158 |
| ON Chronos vs ON control at session 15 | *VirusType-Chronos:Stim+ VirusType-Chronos:Stim:SessionNum\*15* | 0.8 |
| ON Chronos vs OFF Chronos at session 15 | *Stim + VirusType-Chronos:Stim + VirusType-Chronos:Stim:SessionNum\*15* | 0.6837 |
| ON control vs OFF control at session 15 | *Stim + Stim:SessionNum\*15* | 0.6203 |

**Table 3:**
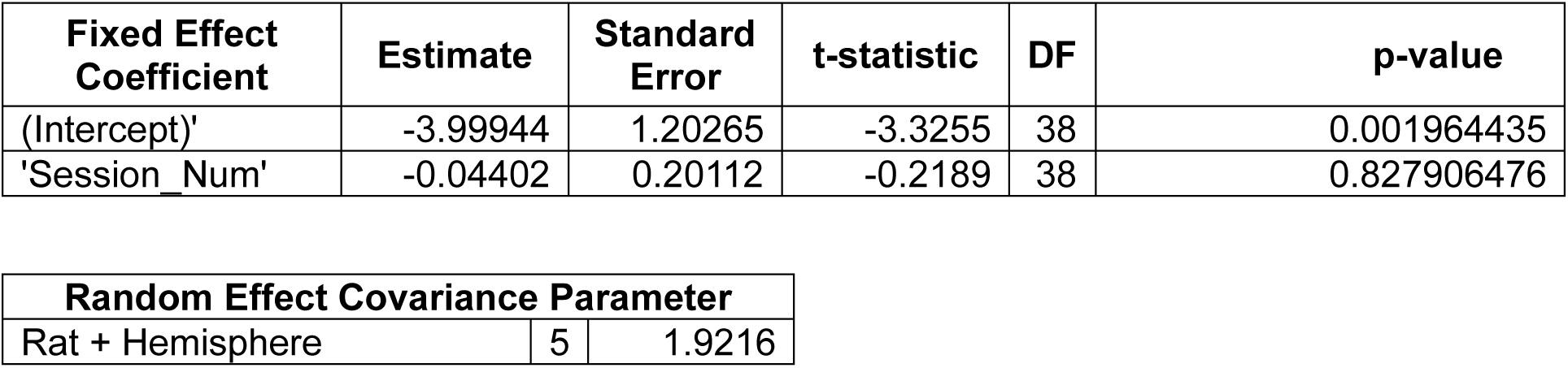
Generalized-linear-model coefficients for the post – pre opto-DBS difference in AUC of z-scored midSTR evoked response potential. Regression coefficients for AUC change after 30 minutes of DBS in z-scored amplitude in the evoked-response potentials experiment, from a GLM with a normal distribution and identity link function. Formula: *AUC_Difference ∼ 1 + SessionNum + (1|Rat_Hemi)*

**Table 4:** Generalized-linear-model coefficients for the post – pre opto-DBS difference in AUC slope of z-scored midSTR evoked response potential.

| Fixed Effect Coefficient | Estimate | Standard Error | t-statistic | DF | p-value |
| --- | --- | --- | --- | --- | --- |
| '(Intercept)' | -0.00040 | 0.00022 | -1.8054 | 38 | 0.078944856 |
| 'Session_Num' | -0.00005 | 0.00005 | -0.8692 | 38 | 0.390173029 |

| Random Effect Covariance Parameter |  |  |
| --- | --- | --- |
| Rat + Hemisphere | 5 | 3.3354E-07 |

**Table 5:**
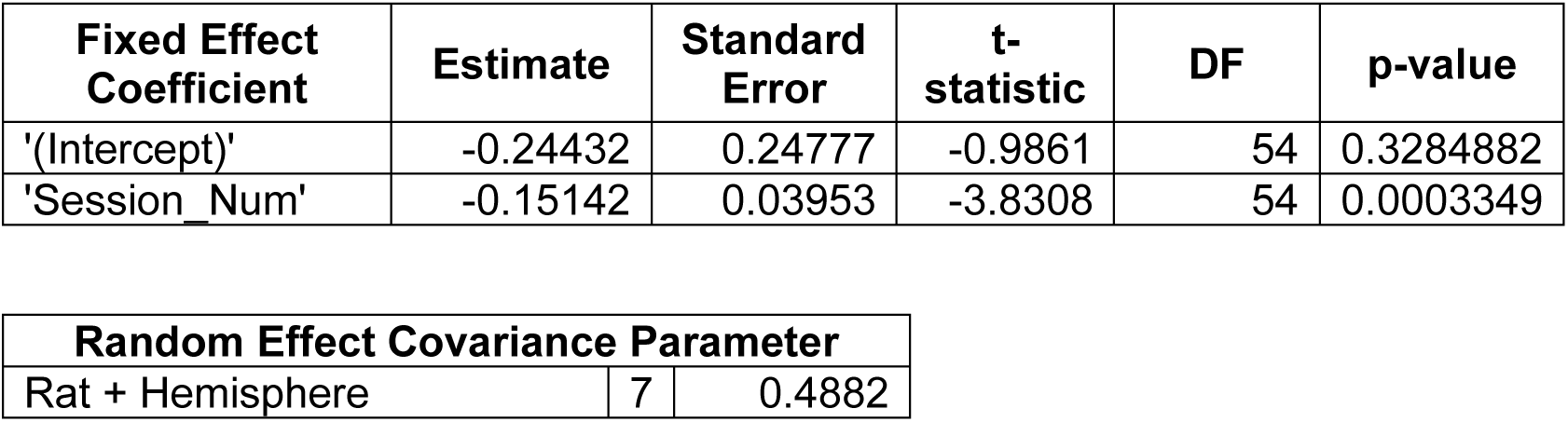
Generalized-linear-model coefficients for the post – pre opto-DBS difference in AUC of z-scored mPFC evoked response potential, early component (6-12 ms after stimulation onset)

**Table 6:** Generalized-linear-model coefficients for the post – pre opto-DBS difference in AUC slope of z-scored mPFC evoked response potential, early component (6-12 ms after stimulation onset)

| Fixed Effect Coefficient | Estimate | Standard Error | t-statistic | DF | p-value |
| --- | --- | --- | --- | --- | --- |
| '(Intercept)' | -0.00005 | 0.00006 | -0.8069 | 54 | 0.423249183 |
| 'Session_Num' | -0.00004 | 0.00001 | -3.7644 | 54 | 0.000413513 |

| Random Effect Covariance Parameter |  |  |
| --- | --- | --- |
| Rat + Hemisphere | 7 | 1.0502E-04 |

**Table 7:** Generalized-linear-model coefficients for the post – pre opto-DBS difference in AUC of z-scored mPFC evoked response potential, late component (0-35 ms after end time of early component)

| Fixed Effect Coefficient | Estimate | Standard Error | t-statistic | DF | p-value |
| --- | --- | --- | --- | --- | --- |
| '(Intercept)' | -1.66331 | 1.19894 | -1.3873 | 54 | 0.171046012 |
| 'Session_Num' | -0.34222 | 0.16175 | -2.1157 | 54 | 0.038997959 |

| Random Effect Covariance Parameter |  |  |
| --- | --- | --- |
| Rat + Hemisphere | 7 | 2.6186 |

**Table 8:** Generalized-linear-model coefficients for the post – pre opto-DBS difference in AUC slope of z-scored mPFC evoked response potential, late component (0-35 ms after end time of early component)

| Fixed Effect Coefficient | Estimate | Standard Error | t-statistic | DF | p-value |
| --- | --- | --- | --- | --- | --- |
| '(Intercept)' | -0.00052 | 0.00034 | -1.5411 | 54 | 0.129139226 |
| 'Session_Num' | -0.00002 | 0.00005 | -0.4355 | 54 | 0.664967807 |

| Random Effect Covariance Parameter |  |  |
| --- | --- | --- |
| Rat + Hemisphere | 7 | 9.4285E-04 |

**Table 9:** Generalized-linear-model coefficients for Set-Shift accuracy in mPFC group.

| Fixed Effect Coefficient | Estimate | Standard Error | t-statistic | DF | p-value |
| --- | --- | --- | --- | --- | --- |
| '(Intercept)' | 0.77762 | 0.06828 | 11.3882 | 3463<sub>9</sub> | 5.41E-30 |
| 'Stim' | 0.05590 | 0.09845 | 0.5678 | 3463<sub>9</sub> | 0.57017284<sub>8</sub> |
| 'SessionNum' | 0.00814 | 0.00386 | 2.1079 | 3463<sub>9</sub> | 0.03504403<sub>4</sub> |
| 'SLRuleInv' | -0.64316 | 0.02434 | -26.4283 | 3463<sub>9</sub> | 2.11E-152 |
| 'Bin_2' | 0.14653 | 0.07210 | 2.0323 | 3463<sub>9</sub> | 0.04212921<sub>7</sub> |
| 'Bin_3' | 0.11440 | 0.07150 | 1.6000 | 3463<sub>9</sub> | 0.10960066<sub>6</sub> |
| 'Bin_4' | 0.14679 | 0.07229 | 2.0305 | 3463<sub>9</sub> | 0.04231500<sub>7</sub> |
| 'Bin_5' | 0.14173 | 0.07224 | 1.9619 | 3463<sub>9</sub> | 0.04977717<sub>7</sub> |
| 'Bin_6' | 0.25379 | 0.07232 | 3.5092 | 3463<sub>9</sub> | 0.00045010<sub>1</sub> |
| 'Bin_7' | 0.19308 | 0.07244 | 2.6654 | 3463<sub>9</sub> | 0.00769343<sub>5</sub> |
| 'Bin_8' | 0.17518 | 0.07194 | 2.4350 | 3463<sub>9</sub> | 0.01489713<sub>5</sub> |
| 'Bin_9' | 0.23440 | 0.07356 | 3.1866 | 3463<sub>9</sub> | 0.00144105<sub>2</sub> |
| 'Bin_10' | 0.30645 | 0.07840 | 3.9090 | 3463<sub>9</sub> | 9.29E-05 |
| 'VirusSite_mPFC-Chronos:Stim' | 0.07864 | 0.11821 | 0.6653 | 3463<sub>9</sub> | 0.50588219<sub>4</sub> |
| 'Stim:SessionNum' | -0.00344 | 0.00673 | -0.5117 | 3463<sub>9</sub> | 0.60883379<sub>1</sub> |
| 'Stim:Bin_2' | -0.05411 | 0.12087 | -0.4476 | 3463<sub>9</sub> | 0.65441033<sub>3</sub> |
| 'Stim:Bin_3' | 0.03079 | 0.12093 | 0.2546 | 3463<sub>9</sub> | 0.79904913<sub>8</sub> |
| 'Stim:Bin_4' | 0.05568 | 0.12069 | 0.4614 | 3463<sub>9</sub> | 0.64453569<sub>6</sub> |
| 'Stim:Bin_5' | 0.09505 | 0.12218 | 0.7780 | 3463<sub>9</sub> | 0.43658758<sub>5</sub> |
| 'Stim:Bin_6' | 0.07312 | 0.12479 | 0.5860 | 3463<sub>9</sub> | 0.55790983<sub>6</sub> |
| 'Stim:Bin_7' | 0.11389 | 0.12558 | 0.9069 | 3463<sub>9</sub> | 0.36446016<sub>9</sub> |
| 'Stim:Bin_8' | 0.08055 | 0.12733 | 0.6326 | 3463<sub>9</sub> | 0.52699264<sub>6</sub> |
| 'Stim:Bin_9' | 0.04793 | 0.12290 | 0.3900 | 3463<sub>9</sub> | 0.69654079<sub>9</sub> |
| 'Stim:Bin_10' | -0.38101 | 0.13079 | -2.9132 | 3463<sub>9</sub> | 0.00357975<sub>3</sub> |
| 'VirusSite_mPFC-Chronos:Stim:SessionNum' | -0.00557 | 0.00757 | -0.7359 | 3463<sub>9</sub> | 0.46177047<sub>3</sub> |
| 'VirusSite_mPFC-Chronos:Stim:Bin_2' | -0.14294 | 0.14157 | -1.0097 | 3463<br>9 | 0.31265765 |
| 'VirusSite_mPFC-Chronos:Stim:Bin_3' | -0.13168 | 0.14298 | -0.9210 | 3463<br>9 | 0.35704605 |
| 'VirusSite_mPFC-Chronos:Stim:Bin_4' | 0.03806 | 0.14309 | 0.2660 | 3463<br>9 | 0.79025639<br>8 |
| 'VirusSite_mPFC-Chronos:Stim:Bin_5' | -0.05920 | 0.14307 | -0.4137 | 3463<br>9 | 0.67906897<br>1 |
| 'VirusSite_mPFC-Chronos:Stim:Bin_6' | -0.08807 | 0.14513 | -0.6068 | 3463<br>9 | 0.54397368<br>8 |
| 'VirusSite_mPFC-Chronos:Stim:Bin_7' | -0.10753 | 0.14438 | -0.7447 | 3463<br>9 | 0.45643149 |
| 'VirusSite_mPFC-Chronos:Stim:Bin_8' | -0.15128 | 0.14611 | -1.0354 | 3463<br>9 | 0.30050486<br>9 |
| 'VirusSite_mPFC-Chronos:Stim:Bin_9' | -0.11205 | 0.14429 | -0.7766 | 3463<br>9 | 0.43740445<br>1 |
| 'VirusSite_mPFC-Chronos:Stim:Bin_10' | 0.30792 | 0.14522 | 2.1203 | 3463<br>9 | 0.03398467<br>6 |

| Random Effects Covariance Parameters |  |  |
| --- | --- | --- |
| Group | Levels | Estimate |
| Chamber | 3 | 3.83E-05 |
| SessionID | 254 | 0.07275 |
| Subject | 16 | 0.11779 |

**Table 10:** Generalized-linear model coefficients for Set-Shift trial omissions in mPFC group.

| Fixed Effect Coefficient | Estimate | Standard Error | t-statistic | DF | p-value |
| --- | --- | --- | --- | --- | --- |
| '(Intercept)' | -4.7301 | 0.3347 | -14.134 | 3549<sub>9</sub> | 3.13E-45 |
| 'Stim' | -0.4609 | 0.6042 | -0.763 | 3549<sub>9</sub> | 0.44554459<sub>9</sub> |
| 'SessionNum' | -0.0516 | 0.0265 | -1.949 | 3549<sub>9</sub> | 0.05127666<sub>5</sub> |
| 'SLRuleInv' | 0.2743 | 0.0820 | 3.347 | 3549<sub>9</sub> | 0.00081731<sub>3</sub> |
| 'Bin_2' | 0.0767 | 0.3359 | 0.228 | 3549<sub>9</sub> | 0.81946092<sub>5</sub> |
| 'Bin_3' | -0.3499 | 0.3718 | -0.941 | 3549<sub>9</sub> | 0.34669651<sub>6</sub> |
| 'Bin_4' | -0.2315 | 0.3651 | -0.634 | 3549<sub>9</sub> | 0.52606788<sub>7</sub> |
| 'Bin_5' | 0.2148 | 0.3297 | 0.652 | 3549<sub>9</sub> | 0.51472780<sub>2</sub> |
| 'Bin_6' | 0.1031 | 0.3365 | 0.306 | 3549<sub>9</sub> | 0.75936881<sub>7</sub> |
| 'Bin_7' | 0.5366 | 0.3078 | 1.743 | 3549<sub>9</sub> | 0.08131578<sub>4</sub> |
| 'Bin_8' | 0.7709 | 0.2962 | 2.603 | 3549<sub>9</sub> | 0.00925176<sub>4</sub> |
| 'Bin_9' | 1.2398 | 0.2758 | 4.495 | 3549<sub>9</sub> | 6.99E-06 |
| 'Bin_10' | 1.3123 | 0.2747 | 4.778 | 3549<sub>9</sub> | 1.78E-06 |
| 'VirusSite_mPFC-Chronos:Stim' | 0.6444 | 0.6385 | 1.009 | 3549<sub>9</sub> | 0.31285233<sub>6</sub> |
| 'Stim:SessionNum' | -0.0130 | 0.0501 | -0.259 | 3549<sub>9</sub> | 0.79575834<sub>8</sub> |
| 'Stim:Bin_2' | 0.2962 | 0.6317 | 0.469 | 3549<sub>9</sub> | 0.63919076<sub>5</sub> |
| 'Stim:Bin_3' | 0.1586 | 0.7233 | 0.219 | 3549<sub>9</sub> | 0.82645134<sub>7</sub> |
| 'Stim:Bin_4' | 0.4134 | 0.6590 | 0.627 | 3549<sub>9</sub> | 0.53042729<sub>7</sub> |
| 'Stim:Bin_5' | -0.6446 | 0.7407 | -0.870 | 3549<sub>9</sub> | 0.38415867<sub>4</sub> |
| 'Stim:Bin_6' | 0.3743 | 0.6322 | 0.592 | 3549<sub>9</sub> | 0.55383742<sub>9</sub> |
| 'Stim:Bin_7' | -0.0256 | 0.6182 | -0.041 | 3549<sub>9</sub> | 0.96703377<sub>6</sub> |
| 'Stim:Bin_8' | 0.0405 | 0.5889 | 0.069 | 3549<sub>9</sub> | 0.94514218<sub>2</sub> |
| 'Stim:Bin_9' | -0.5548 | 0.5622 | -0.987 | 3549<sub>9</sub> | 0.32372391<sub>5</sub> |
| 'Stim:Bin_10' | 0.0758 | 0.5355 | 0.142 | 3549<sub>9</sub> | 0.88747372<sub>3</sub> |
| 'VirusSite_mPFC-Chronos:Stim:SessionNum' | 0.0582 | 0.0543 | 1.072 | 3549<br>9 | 0.28387192<br>8 |
| 'VirusSite_mPFC-Chronos:Stim:Bin_2' | -0.8230 | 0.6822 | -1.206 | 3549<br>9 | 0.22770417<br>1 |
| 'VirusSite_mPFC-Chronos:Stim:Bin_3' | -0.1679 | 0.7452 | -0.225 | 3549<br>9 | 0.82175448<br>9 |
| 'VirusSite_mPFC-Chronos:Stim:Bin_4' | -1.0418 | 0.7250 | -1.437 | 3549<br>9 | 0.15074243<br>9 |
| 'VirusSite_mPFC-Chronos:Stim:Bin_5' | -0.1763 | 0.7946 | -0.222 | 3549<br>9 | 0.82444796<br>1 |
| 'VirusSite_mPFC-Chronos:Stim:Bin_6' | -0.6338 | 0.6608 | -0.959 | 3549<br>9 | 0.33746384<br>9 |
| 'VirusSite_mPFC-Chronos:Stim:Bin_7' | -0.4009 | 0.6441 | -0.622 | 3549<br>9 | 0.53363053<br>6 |
| 'VirusSite_mPFC-Chronos:Stim:Bin_8' | -0.2432 | 0.6068 | -0.401 | 3549<br>9 | 0.68861166 |
| 'VirusSite_mPFC-Chronos:Stim:Bin_9' | 0.4114 | 0.5828 | 0.706 | 3549<br>9 | 0.48029936<br>9 |
| 'VirusSite_mPFC-Chronos:Stim:Bin_10' | 0.2662 | 0.5492 | 0.485 | 3549<br>9 | 0.62779777<br>9 |

| Random Effects Covariance Parameters |  |  |
| --- | --- | --- |
| Group | Levels | Estimate |
| Chamber | 3 | 2.09E-06 |
| SessionID | 254 | 1.0309 |
| Subject | 16 | 2.25E-13 |

**Table 11:** Generalized-linear model coefficients for Set-Shift accuracy in midSTR group.

| Fixed Effect Coefficient | Estimate | Standard Error | t-statistic | DF | p-value |
| --- | --- | --- | --- | --- | --- |
| '(Intercept)' | 0.8847 | 0.0676 | 13.091 | 4123<br>2 | 4.44E-39 |
| 'Stim' | -0.0031 | 0.0948 | -0.033 | 4123<br>2 | 0.9739569<br>6 |
| 'SessionNum' | 0.0038 | 0.0032 | 1.198 | 4123<br>2 | 0.2310192<br>9 |
| 'SLRuleInv' | -0.6766 | 0.0220 | -30.723 | 4123<br>2 | 5.82E-205 |
| 'Bin_2' | 0.0914 | 0.0653 | 1.400 | 4123<br>2 | 0.1616041<br>3 |
| 'Bin_3' | 0.0265 | 0.0652 | 0.406 | 4123<br>2 | 0.6847749<br>5 |
| 'Bin_4' | 0.0530 | 0.0656 | 0.808 | 4123<br>2 | 0.4190059<br>5 |
| 'Bin_5' | 0.0944 | 0.0655 | 1.440 | 4123<br>2 | 0.1498287<br>1 |
| 'Bin_6' | 0.1400 | 0.0658 | 2.129 | 4123<br>2 | 0.0332957<br>7 |
| 'Bin_7' | 0.1338 | 0.0658 | 2.032 | 4123<br>2 | 0.0421585<br>5 |
| 'Bin_8' | 0.0976 | 0.0650 | 1.500 | 4123<br>2 | 0.1336312<br>2 |
| 'Bin_9' | 0.1852 | 0.0637 | 2.905 | 4123<br>2 | 0.0036689 |
| 'Bin_10' | 0.3079 | 0.0657 | 4.683 | 4123<br>2 | 2.84E-06 |
| 'VirusSite_midSTR-Chronos:Stim' | -0.1410 | 0.1086 | -1.298 | 4123<br>2 | 0.1944326<br>4 |
| 'Stim:SessionNum' | 0.0089 | 0.0058 | 1.530 | 4123<br>2 | 0.1259533<br>8 |
| 'Stim:Bin_2' | -0.0623 | 0.1155 | -0.540 | 4123<br>2 | 0.5894471<br>8 |
| 'Stim:Bin_3' | -0.0116 | 0.1161 | -0.100 | 4123<br>2 | 0.9204044<br>3 |
| 'Stim:Bin_4' | -0.0138 | 0.1163 | -0.118 | 4123<br>2 | 0.9057690<br>8 |
| 'Stim:Bin_5' | -0.1846 | 0.1160 | -1.591 | 4123<br>2 | 0.11160880 |
| 'Stim:Bin_6' | -0.1039 | 0.1163 | -0.893 | 4123<br>2 | 0.3717758<br>8 |
| 'Stim:Bin_7' | 0.0387 | 0.1171 | 0.330 | 4123<br>2 | 0.7410995<br>7 |
| 'Stim:Bin_8' | 0.0098 | 0.1183 | 0.082 | 4123<br>2 | 0.9343015<br>6 |
| 'Stim:Bin_9' | -0.0826 | 0.1206 | -0.685 | 4123<br>2 | 0.4932322<br>3 |
| 'Stim:Bin_10' | -0.3426 | 0.1134 | -3.021 | 4123<br>2 | 0.0025210<br>3 |
| 'VirusSite_midSTR-Chronos:Stim:SessionNum' | 0.0019 | 0.0067 | 0.288 | 4123<br>2 | 0.7730368<br>4 |
| 'VirusSite_midSTR-Chronos:Stim:Bin_2' | 0.1422 | 0.1316 | 1.080 | 4123<br>2 | 0.2800074<br>5 |
| 'VirusSite_midSTR-Chronos:Stim:Bin_3' | 0.1734 | 0.1318 | 1.315 | 4123<br>2 | 0.1883677<br>2 |
| 'VirusSite_midSTR-Chronos:Stim:Bin_4' | 0.0726 | 0.1314 | 0.552 | 4123<br>2 | 0.5806836<br>2 |
| 'VirusSite_midSTR-Chronos:Stim:Bin_5' | 0.2693 | 0.1316 | 2.047 | 4123<br>2 | 0.0406972<br>5 |
| 'VirusSite_midSTR-Chronos:Stim:Bin_6' | 0.1620 | 0.1319 | 1.228 | 4123<br>2 | 0.2192972<br>6 |
| 'VirusSite_midSTR-Chronos:Stim:Bin_7' | -0.0281 | 0.1327 | -0.212 | 4123<br>2 | 0.8324508<br>3 |
| 'VirusSite_midSTR-Chronos:Stim:Bin_8' | 0.1377 | 0.1346 | 1.023 | 4123<br>2 | 0.3062146<br>0 |
| 'VirusSite_midSTR-Chronos:Stim:Bin_9' | 0.1529 | 0.1407 | 1.087 | 4123<br>2 | 0.2769761<br>5 |
| 'VirusSite_midSTR-Chronos:Stim:Bin_10' | 0.2428 | 0.1411 | 1.721 | 4123<br>2 | 0.0852093<br>7 |

| Random Effects Covariance Parameters |  |  |
| --- | --- | --- |
| Group | Levels | Estimate |
| Chamber | 4 | 6.50E-02 |
| SessionID | 304 | 0.022479 |
| Subject | 19 | 0.09917 |

**Table 12:**
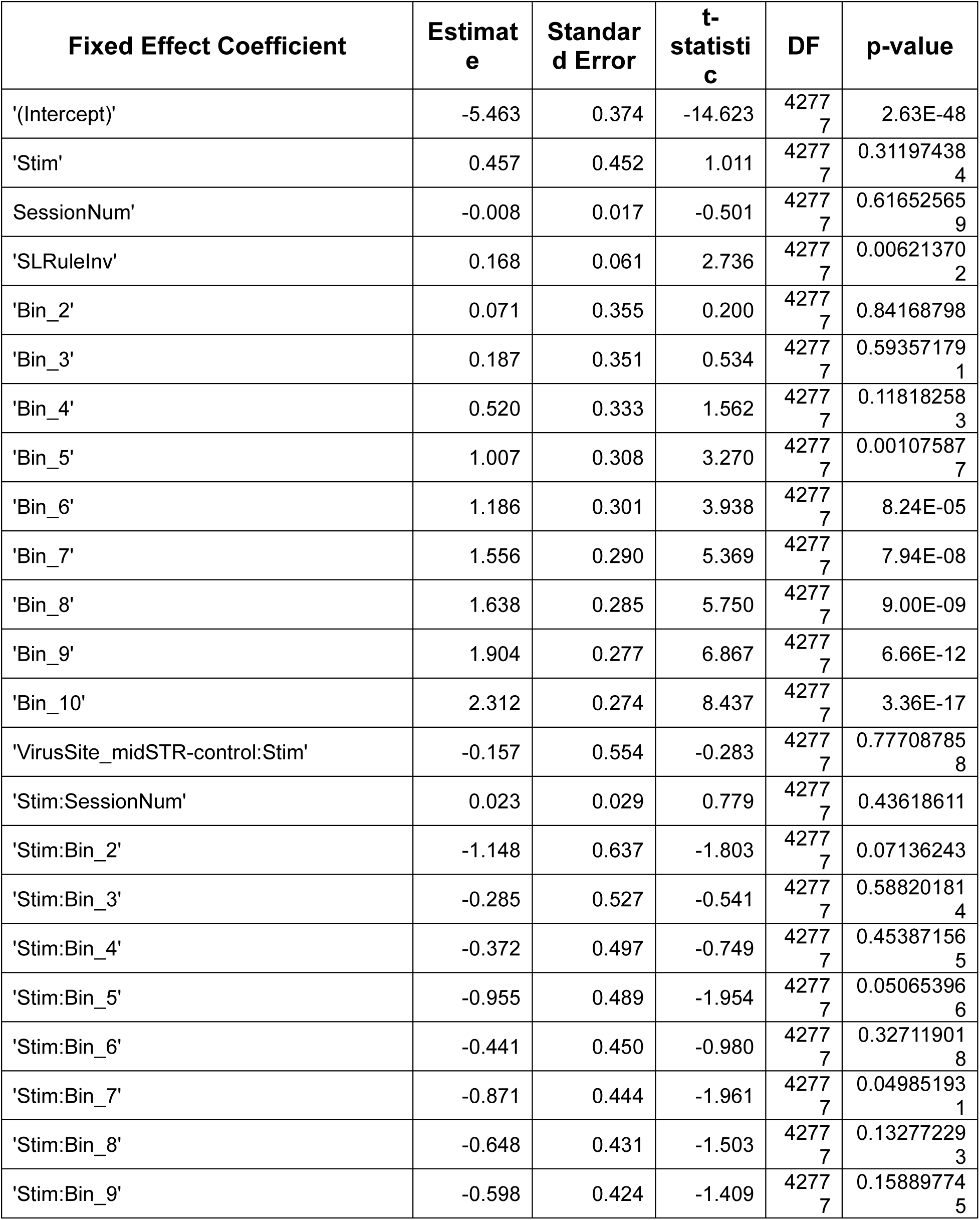
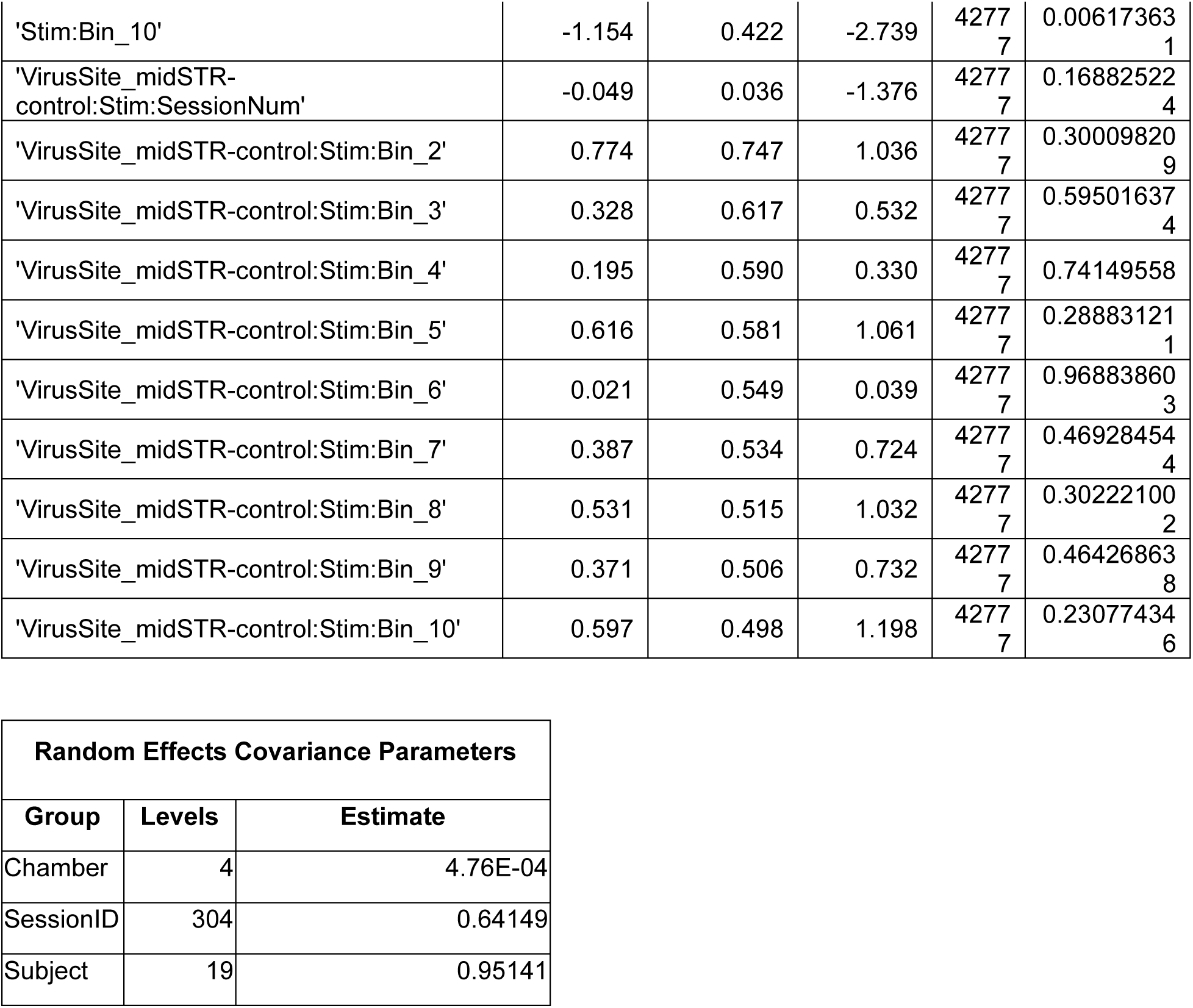
Generalized-linear model coefficients for Set-Shift trial omissions in midSTR group.

| Fixed Effect Coefficient | Estimate | Standard Error | t-statistic | DF | p-value |
| --- | --- | --- | --- | --- | --- |
| '(Intercept)' | -5.463 | 0.374 | -14.623 | 4277<br>7 | 2.63E-48 |
| 'Stim' | 0.457 | 0.452 | 1.011 | 4277<br>7 | 0.31197438<br>4 |
| SessionNum' | -0.008 | 0.017 | -0.501 | 4277<br>7 | 0.61652565<br>9 |
| 'SLRuleInv' | 0.168 | 0.061 | 2.736 | 4277<br>7 | 0.00621370<br>2 |
| 'Bin_2' | 0.071 | 0.355 | 0.200 | 4277<br>7 | 0.84168798 |
| 'Bin_3' | 0.187 | 0.351 | 0.534 | 4277<br>7 | 0.59357179<br>1 |
| 'Bin_4' | 0.520 | 0.333 | 1.562 | 4277<br>7 | 0.11818258<br>3 |
| 'Bin_5' | 1.007 | 0.308 | 3.270 | 4277<br>7 | 0.00107587<br>7 |
| 'Bin_6' | 1.186 | 0.301 | 3.938 | 4277<br>7 | 8.24E-05 |
| 'Bin_7' | 1.556 | 0.290 | 5.369 | 4277<br>7 | 7.94E-08 |
| 'Bin_8' | 1.638 | 0.285 | 5.750 | 4277<br>7 | 9.00E-09 |
| 'Bin_9' | 1.904 | 0.277 | 6.867 | 4277<br>7 | 6.66E-12 |
| 'Bin_10' | 2.312 | 0.274 | 8.437 | 4277<br>7 | 3.36E-17 |
| 'VirusSite_midSTR-control:Stim' | -0.157 | 0.554 | -0.283 | 4277<br>7 | 0.77708785<br>8 |
| 'Stim:SessionNum' | 0.023 | 0.029 | 0.779 | 4277<br>7 | 0.43618611 |
| 'Stim:Bin_2' | -1.148 | 0.637 | -1.803 | 4277<br>7 | 0.07136243 |
| 'Stim:Bin_3' | -0.285 | 0.527 | -0.541 | 4277<br>7 | 0.58820181<br>4 |
| 'Stim:Bin_4' | -0.372 | 0.497 | -0.749 | 4277<br>7 | 0.45387156<br>5 |
| 'Stim:Bin_5' | -0.955 | 0.489 | -1.954 | 4277<br>7 | 0.05065396<br>6 |
| 'Stim:Bin_6' | -0.441 | 0.450 | -0.980 | 4277<br>7 | 0.32711901<br>8 |
| 'Stim:Bin_7' | -0.871 | 0.444 | -1.961 | 4277<br>7 | 0.04985193<br>1 |
| 'Stim:Bin_8' | -0.648 | 0.431 | -1.503 | 4277<br>7 | 0.13277229<br>3 |
| 'Stim:Bin_9' | -0.598 | 0.424 | -1.409 | 4277<br>7 | 0.15889774<br>5 |
| 'Stim:Bin_10' | -1.154 | 0.422 | -2.739 | 4277<br>7 | 0.00617363<br>1 |
| 'VirusSite_midSTR-control:Stim:SessionNum' | -0.049 | 0.036 | -1.376 | 4277<br>7 | 0.16882522<br>4 |
| 'VirusSite_midSTR-control:Stim:Bin_2' | 0.774 | 0.747 | 1.036 | 4277<br>7 | 0.30009820<br>9 |
| 'VirusSite_midSTR-control:Stim:Bin_3' | 0.328 | 0.617 | 0.532 | 4277<br>7 | 0.59501637<br>4 |
| 'VirusSite_midSTR-control:Stim:Bin_4' | 0.195 | 0.590 | 0.330 | 4277<br>7 | 0.74149558 |
| 'VirusSite_midSTR-control:Stim:Bin_5' | 0.616 | 0.581 | 1.061 | 4277<br>7 | 0.28883121<br>1 |
| 'VirusSite_midSTR-control:Stim:Bin_6' | 0.021 | 0.549 | 0.039 | 4277<br>7 | 0.96883860<br>3 |
| 'VirusSite_midSTR-control:Stim:Bin_7' | 0.387 | 0.534 | 0.724 | 4277<br>7 | 0.46928454<br>4 |
| 'VirusSite_midSTR-control:Stim:Bin_8' | 0.531 | 0.515 | 1.032 | 4277<br>7 | 0.30222100<br>2 |
| 'VirusSite_midSTR-control:Stim:Bin_9' | 0.371 | 0.506 | 0.732 | 4277<br>7 | 0.46426863<br>8 |
| 'VirusSite_midSTR-control:Stim:Bin_10' | 0.597 | 0.498 | 1.198 | 4277<br>7 | 0.23077434<br>6 |

| Random Effects Covariance Parameters |  |  |
| --- | --- | --- |
| Group | Levels | Estimate |
| Chamber | 4 | 4.76E-04 |
| SessionID | 304 | 0.64149 |
| Subject | 19 | 0.95141 |

**Table 13:** GLM comparison for different bin sizes for mPFC group.

| Number of Bins | Number of Fixed Effects | Model AIC |
| --- | --- | --- |
| 1 | 7 | 21109 |
| 2 | 10 | 19381 |
| 3 | 13 | 18890 |
| 4 | 16 | 18603 |
| 5 | 19 | 18373 |
| 6 | 22 | 18306 |
| 7 | 25 | 18254 |
| 8 | 28 | 18172 |
| 9 | 31 | 18146 |
| 10 | 34 | 18151 |
| 11 | 37 | 18111 |
| 12 | 40 | 18112 |
| 13 | 43 | 18093 |
| 14 | 46 | 18099 |
| 15 | 49 | 18075 |

**Table 14:**
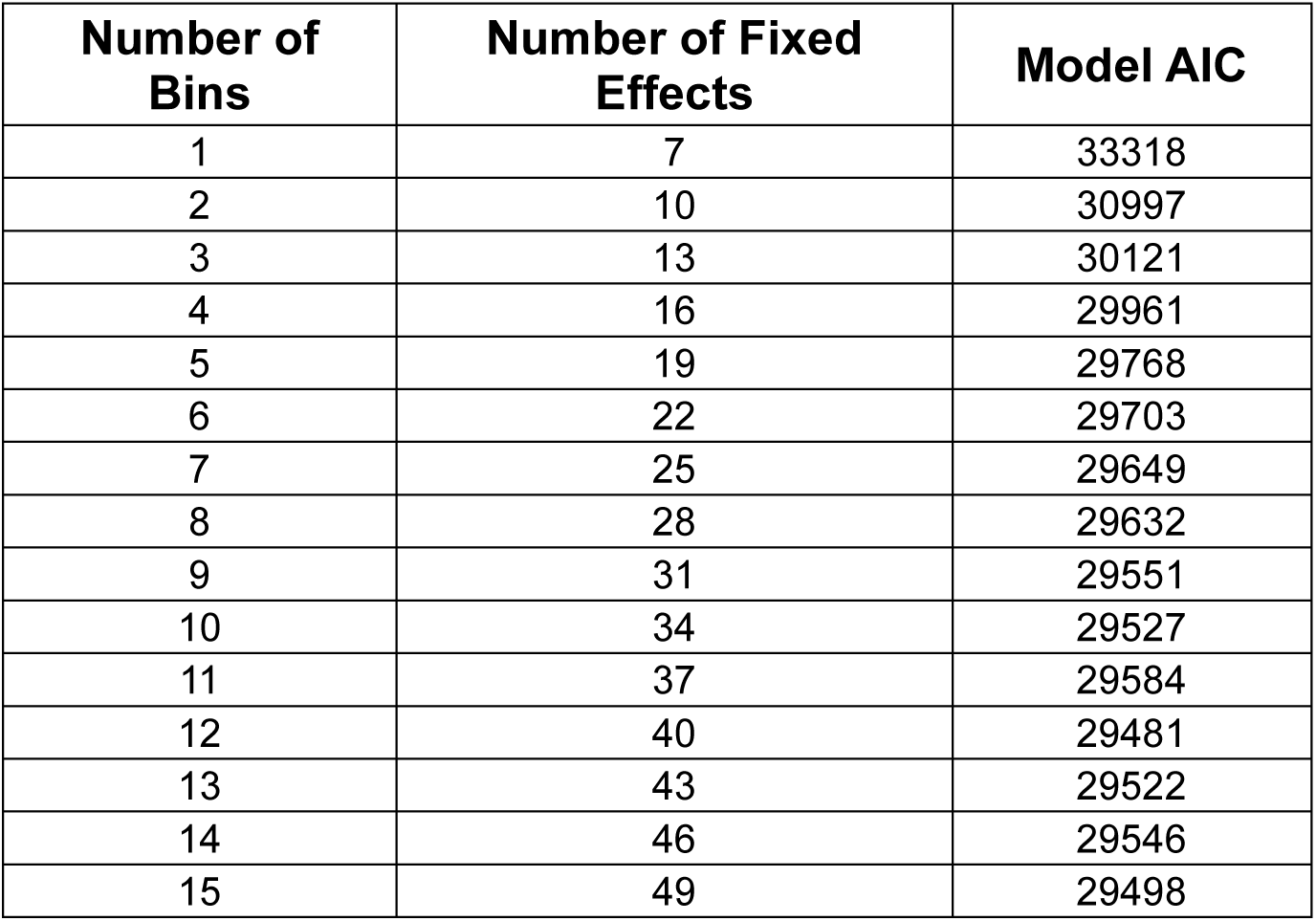
GLM comparison for different bin sizes for midSTR group.

**Table 15:** Generalized Linear Model Coefficients for RT in midSTR electrical DBS study (adapted from [8]) Regression coefficients for response time (RT) in seconds in the mPFC-axon opto-DBS experiment from a generalized linear mixed-effects model (GLM), with a gamma distribution and identity link function. (Here, we use raw RespTime instead of binned time-in-session, but they effectively mean the same thing). **Formula:** *RT ∼ 1 + RuleType + Stim*SessionNum + Stim*RespTime + (1 | Rat) + (1 | Session)*

| Fixed Effect Coefficient | Estimate | Standard Error | t-statistic | DF | p-value |
| --- | --- | --- | --- | --- | --- |
| '(Intercept)' | 0.70091 | 0.04810 | 14.5720 | 17815 | 7.94E-48 |
| Stim_ON' | -0.08152 | 0.03681 | -2.2147 | 17815 | 0.026792 |
| SessionNum' | -0.00779 | 0.00216 | -3.6102 | 17815 | 0.0003068 |
| TimeInSession' | 0.00004 | 6.62170E-06 | 6.0780 | 17815 | 1.24E-09 |
| SLRule_1' | 0.01059 | 0.00386 | 2.7431 | 17815 | 0.0060917 |
| Stim_1ON:SessionNum' | 0.00430 | 0.00341 | 1.2600 | 17815 | 0.20767 |
| Stim_ON:RespTime' | -1.69550E-06 | 9.20680E-06 | -0.1842 | 17815 | 0.85389 |

| Random Effects Covariance Parameters |  |  |
| --- | --- | --- |
| Group | Levels | Estimate |
| Rat | 8 | 0.12063 |
| SessionID | 134 | 0.092948 |

### Statistical analysis

#### Design and statistical power

For Set-Shift opto-DBS experiments, we pre-determined sample size and number of sessions based on prior work [8] and a power analysis (G*Power 3.1). A generalized-linear model (GLM) is functionally equivalent to a repeated-measures ANOVA, with within-between interactions, as we are comparing opto-DBS ON vs. OFF effects in both Chronos and control rats. To achieve 80% power with an effect size of f = 0.25 and a Bonferroni-corrected alpha = 0.0167, we required 8 animals minimum per group. We therefore targeted an N = 8 to 10 per group, balancing numbers in accordance with pair-housing rats.

#### Statistical analysis of Set-Shift behavior

Consistent with prior work [8,27] generalized-linear models (GLMs) were generated in MATLAB (R2023b) for behavioral analysis of RT, accuracy, and omissions. GLMs can effectively handle repeated-measures data (different stimulation conditions applied to the same animal) and the non-Gaussian distribution of RTs. GLMs were designed to closely follow those used in electrical stimulation studies. We included fixed effects for Stimulation (ON or OFF), SessionNumber (1 to 16, total number of testing sessions), RuleType (Side or Light), and binned Time-in-session (TIS).

The TIS is the time in a session a rat responded (made a choice) on a Set-Shift trial. To measure how RT varied with the TIS and its interactions with Stim and VirusType, we binned the 0 to 90 min range of TISs, with each bin containing an equal number of TISs. We compared the AICs from GLMs with different numbers of bins ranging from 1 to 15. For mPFC rats, 10 bins resulted in the lowest AIC while maintaining an adequate number of TISs in each bin for statistics and plotting. For midSTR rats, 5 and 10 bins resulted in the lowest AICs, so 10 bins was chosen to maintain consistency with the mPFC experiment and for visual comparison between the two groups (Figures 2D and 3D).

In addition to these fixed effects, we added key interaction terms, such as Stimulation and VirusType (Chronos or control), and Stimulation and SessionNumber, as well as random effect terms for Subject, Chamber, and SessionID. P-values for GLM coefficients and combinations of terms (e.g. Opto-DBS ON RTs in Chronos compared to control rats at bin 2), were computed using the *coefTest* MATLAB function (see post-hoc comparisons in Supplementary Tables). The value of SessionID ranged from 1 to ∼300 (16 sessions * total number of included rats) and accounted for correlation among trials from the same session). Omission trials, where the rat did not respond during the 3 s trial window, were excluded from all RT and accuracy analyses. Separate GLMs were used for mPFC-axon opto-DBS vs. midSTR-neuron opto-DBS experiments. The general form of these GLMs is shown below:

*DependentVariable ∼ Stimulation + SessionNumber + RuleType + BinnedTIS + Stimulation:VirusType + Stimulation:BinnedTIS + Stimulation:SessionNumber + Stimulation:VirusType:SessionNumber + Stimulation:VirusType:BinnedTIS + (1|Chamber) + (1|Subject) + (1|SessionID)*

To better visually isolate opto-DBS’s effect on RT, we plotted change in RTs adjusted based on the GLM outputs. The contribution of all fixed and random effects were subtracted from RTs. Then, the effects of interest (e.g. *Stim* and *Stim:VirusType*, Figure 2C) were added back in, leaving just the desired variable(s) impact on RT. These adjusted-RTs were plotted for each group and within and across Set-Shift sessions. .

### Electrophysiology experiment analysis

#### Data Validation and Preprocessing

LFPs were processed offline with custom scripts in MATLAB (2023b). Raw signals were epoched from -2 to 2ms around stimulus onset, down sampled to 2 kHz and filtered to a passband of 4 to 200 Hz (separate high and low pass FIR filters). In each recording site, the channel with the greatest raw ERP amplitude was selected for primary analysis. Due to damage or misplacement during surgical implantation, only unilateral striatal ERPs were present in 3 of 4 rats, and the unresponsive striatal site was excluded from analysis. However, we have previously shown that unilateral electrical midSTR stimulation reproduces bilateral effects [27], and there are contralateral and ipsilateral projections from the mPFC to the striatum [10,45], therefore we included both mPFC sites in the analysis if ERPs were detected (95% confidence intervals of 5V response were separate from 0V sham intervals) during a pre-DBS amplitude sweep session.

#### Evoked Response Potential Analysis

To visualize ERPs, each channel was independently z-scored relative to its 0V (sham) response across timepoints. For each channel and amplitude, pre- and post-DBS pulse trains were rejected from the z-score calculation if response amplitude was greater than 400 µV, a limit determined from the overall noise level, and which resulted in similar numbers of rejected trials in all rats. For the channels selected for further analysis, this resulted in a rejection average of ∼6% of trials for midSTR channels and ∼5% of trials for mPFC channels, per session across rats and pre- and post- tests. Next, z-scores -10 to 200 ms around stimulus onset were baseline-corrected using a baseline of -4.5 to 0 ms for midSTR and -0.5 to 4 ms for mPFC and averaged over trials to obtain the ERP. 95% Confidence intervals were calculated using a non-parametric bootstrap across trials.

To assess ERP dynamics within and across days, we calculated the area under the curve (AUC) by summing midSTR and mPFC responses to the 5V/500µA amplitude (max) stimulation. To determine the AUC time window, we predefined time regions of interest based on stereotypical ERP features in each site observed in this experiment and supported by other studies [46–49]. The midSTR ERP was very consistent across rats, so a uniform region of 0 to 25ms was used to calculate midSTR AUCs for all rats and sites.

We focused on two features of the mPFC ERP, an early negative peak component and a late biphasic component. Because of slight variation in the timing of mPFC responses across rats, we defined early and late components uniquely for each rat + recording site. For the early mPFC component, we calculated AUCs by summing z-scores around the minimum value (peak) found in the early (6 to 12 ms) period of interest ± 3 ms. For the late mPFC component we calculated AUCs by summing values from 0 to 35ms after the end time of the early component. AUC magnitude was used as a readout of the change in circuit strength or output capacity after axonal opto-DBS [50,51]. The post-DBS change in AUC was averaged across all rats+hemisphere for each ERP component and each session (Figure 4C).

**Figure 4:**
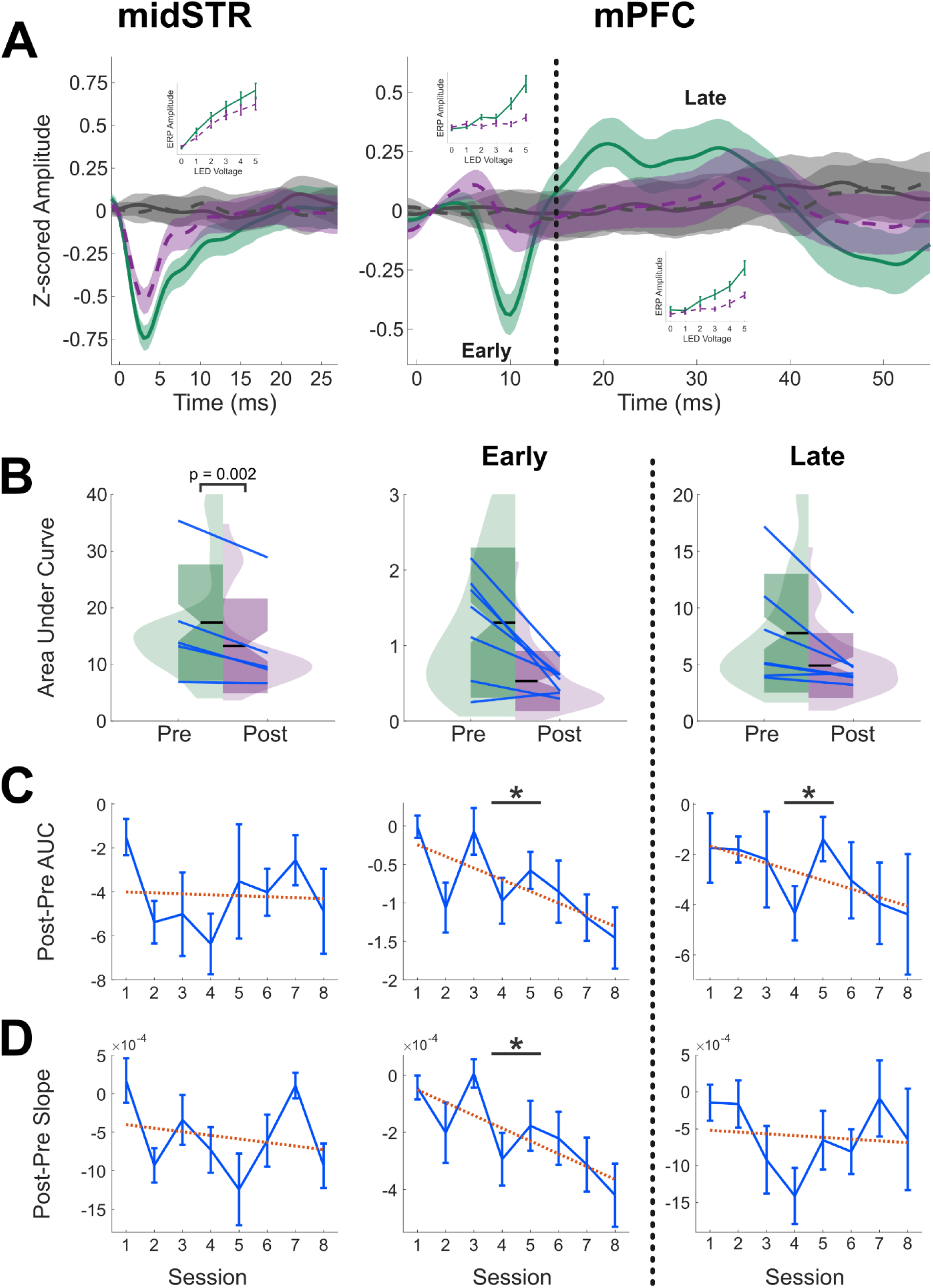
Modulation of evoked-response potentials after mPFC-axon opto-DBS within and across days A) midSTR (left): Example 5V evoked-responses (z-scored amplitude) in midSTR, pre (solid green) and post (dotted purple) 30 min opto-DBS. Gray lines show response to sham (0V) stimulation. Inset plots the mean response curve over 1V to 5V stimulation amplitudes, pre and post DBS. **mPFC (right):** Same as midSTR but epoched into an early and a late response component (vertical gray line shows boundary). **B)** AUC Distributions for each ERP component of interest, across all rats and sessions. Blue lines represent the individual mean of the AUC for each rat+site pre and post opto-DBS. **C)** Post-DBS change in AUC across the 8 test sessions. Blue lines indicate Z-scored AUCs that were averaged across rats+sites at each session. Red dotted line illustrates the slope of the *SessionNum* fixed effect GLM term for that site (* = p < 0.05). **D)** Post-DBS change in the slope of the dose-response curve from 1V to 5V. Blue lines indicate slopes averaged across rats+sites at each session. Red dotted line illustrates the slope of the *SessionNum* fixed effect GLM term for that site (* = p < 0.05).

We also analyzed post-pre changes in the slope of line fit to ERP AUC versus LED input from 1 to 5V (i.e., we generated a dose-response curve, see Figure 4A insets). These curves serve as a metric of the functional strength of the mPFC-midSTR circuit, also referred to as the “gain” of the system [50]. Fit lines indicate the slope of the SessionNum intercept from GLMs (Figure 4D and see below). GLMs for each ERP component were generated to assess the significance of post-DBS changes with increasing session number and included a random effect for each rat+hemisphere. The general format of each GLM was:

*Post - Pre change in AUC/Slope ∼ 1+ SessionNumber + (1 | RatHemi)*

## Results

We used the Set-Shift task to assess opto-DBS’s cognitive control effects (Figure 1A). This task was previously used to develop the preclinical electrical stimulation model of VCVS DBS [8] and in subsequent studies [27,28]. A total of 37 rats, split between 4 groups, were successfully implanted and completed 16 Set-Shift testing sessions, alternating opto-DBS ON or OFF. OFF trial behaviors replicated those in prior work. Rats were less accurate (Figure 1C) and made more omissions (Figure 1D) on Side rule trials compared to Light rule but had similar RTs between rule types (Figure 1B).

### Opto-DBS of mPFC axons improves cognitive control, partially reproducing electrical stimulation effects

Rats assigned to the mPFC group were transfected with Chronos or control - CaMKII-GFP in mPFC subregions and implanted with optic fibers in the midSTR to specifically target cortical axons for opto-DBS during the Set-Shift task. We assessed opto-DBS’s effect on cognitive control by comparing opto-DBS-ON RTs in Chronos vs. control rats, and within-animal ON vs. OFF RTs. ON-Chronos RTs were significantly faster (∼ 100 ms) than ON-Control and OFF-Chronos RTs (Figure 2C), demonstrating improved cognitive control in rats with active DBS, and replicating effects of midSTR electrical stimulation (∼ 50 ms reduction) [8,27]. Control rats were not affected by stimulation (Figure 2C, left). There were no significant changes in accuracy or trial omissions from opto-DBS (Tables S9 and S10). Thus, PFC activation via axons terminating in/passing through mid-striatum is sufficient to produce the previously reported cognitive effect.

Continuous high frequency optogenetic stimulation for sustained periods of time (throughout the duration of the task) is infrequently used in *in vivo* animal models. To address this, we investigated how opto-DBS behavioral effects varied within and across Set-Shift sessions. We isolated the effect of the time in a session when a response occurred (TIS) on RT, for ON vs. OFF and Control vs. Chronos conditions. The RT effect from active axonal opto-DBS declined throughout a session, so that by the 30 min mark, ON-Chronos vs. ON-control rat RTs did not significantly differ (Figure 2D). Similarly, the RT effect declined across sessions so that by the final testing session ON-Chronos and ON-control rat RTs did not significantly differ (Figure 2E). This contrasts with the RT effect from electrical stimulation which remained stable within and across sessions (Table S15).

### Sustained Opto-DBS of local midSTR neurons transiently impairs cognitive control

To test the role of local striatal neurons in DBS’s cognitive control enhancement, a separate group of rats were transfected with Chronos or control -hSyn-GFP at the midSTR stimulation target, alongside optic fibers. Opto-DBS targeted specifically to local striatal neurons had no lasting cognitive control effect. There were no significant differences in ON-Chronos vs. ON-control or OFF-Chronos RTs (Figure 2C). Opto-DBS also had no significant effects on accuracy or trial omissions (Tables S10 and S11). Next, we investigated changes in opto-DBS effects within and across testing days. Within a Set-Shift session, midSTR-neuron opto-DBS starts to impair cognitive control. After ∼15 to 30 min of DBS, ON-Chronos RTs were significantly slower than ON-control (Figure 3D). Across sessions, there was no change in opto-DBS effects (Figure 3E). These results suggest that local striatal neurons are not the main component driving electrical DBS’s effects and may actually impede cognitive control.

### mPFC-axon opto-DBS alters mPFC and midSTR activity within and across days

Interestingly, both mPFC-axon and local neuronal opto-DBS had detrimental effects on cognitive control within a day, but only mPFC-axon opto-DBS effects declined across the 16 Set-Shift sessions. We investigated the possible origins of these temporal changes by comparing evoked-response potentials (ERPs) before and after 30 min of axon opto-DBS across testing days. ERPs can be used to assess how effective a stimulation paradigm is at modulating neural signals or a particular region/circuit [47,48,52]. A separate group of rats (N = 4) were transfected with Chronos in the mPFC and implanted with recording electrodes in the mPFC and in the midSTR alongside optic fibers for stimulation. Rats underwent 8 recording sessions, following the same schedule as behavioral testing. We compared the area-under-the-curve (AUC) of z-scored amplitudes of the local midSTR ERP and an early and a late component of the mPFC ERP, pre and post opto-DBS (Figure 3A). Opto-DBS targeted to cortical axons significantly reduced the magnitude of the local midSTR ERP but did not significantly impact either of the mPFC ERP components (Figure 4B). This provides physiological evidence for the within-day decline in mPFC-axon opto-DBS’s cognitive effects and suggests this change is mediated by striatal synapses.

Next, we assessed opto-DBS-driven changes in mPFC and midSTR circuitry across days. The post-pre change in AUC was computed for each day and averaged across rats. For both mPFC components, pre/post attenuation in AUC from opto-DBS significantly declined across testing days, in other words, the mPFC ERP was attenuated more by opto-DBS with each additional day of testing (Figure 4C, middle and right panels). In contrast, the local midSTR pre/post change remained stable (Figure 4C left panel). To assess how responsive midSTR and mPFC circuit components were pre and post opto-DBS, we calculated the slope of the dose-response curve of different stimulus amplitudes (voltage of LED driver) and the resulting ERP magnitude. There was a significant decrease in the magnitude of the mPFC-early component, e.g. as session number increased, higher stimulus amplitudes were required to achieve the same response magnitude as prior sessions (Figure 4D). The slope change of the midSTR and mPFC-late components were stable over sessions. These results link the decline in axonal opto-DBS’s cognitive enhancement across sessions to changes in how efficiently opto-DBS activates mPFC neurons. The change in the mPFC-early dose-response curve over days suggests a neuroplastic-like mechanism. Overall, these results link the time-dependent changes in mPFC-axon opto-DBS’s cognitive control effects to modulation of specific neural components: within-day changes are reflected by reduced midSTR activity and across-day changes are associated with a decline in mPFC responsiveness.

## Discussion

Here, we employed pathway-specific optogenetic techniques to dissect circuit-level mechanisms underlying the cognitive effects of VCVS DBS. We show that high-frequency stimulation of mPFC axons in the midSTR is sufficient to reproduce the cognitive control enhancement observed with electrical DBS. In contrast, stimulation of local midSTR neurons failed to improve and eventually impaired cognitive control. In addition, we identified time-dependent attenuations in the effect of opto-DBS, where the behavioral benefit of axonal stimulation decayed within a testing session and across days. This “fading” effect correlated with physiological signatures of reduced excitability within testing days in the midSTR and across testing days in the mPFC. Together, these findings suggest that VCVS DBS improves cognitive control by engaging axons descending from the PFC rather than local striatal gray matter nuclei, providing a rationale for targeting white matter tracts in psychiatric DBS therapies.

Our observation that mPFC-axon stimulation drives the cognitive control improvement aligns with recent clinical data suggesting that optimal engagement of white matter tracts may mediate the effect of DBS in OCD and MDD [9,24,53,54]. Mechanistically, high-frequency stimulation of mPFC axons may enhance cognitive control via retrograde/antidromic activation of prefrontal neurons or by coordinating activity (e.g. feedforward inhibition) in the striatum and other downstream regions receiving prefrontal input [55,56]. Set-shifting and other cognitive control functions are linked to both prefrontal and striatal circuits [5,13,57–59]. Specifically, mPFC is thought to serve as a top-down control signal for downstream areas like the striatum [18,60,61]. By engaging this pathway, axonal opto-DBS may effectively increase the gain on these top-down signals, facilitating striatal output and resulting in optimal Set-Shift behavior. Overall, this supports the “network hypothesis” of DBS, positing that its therapeutic effects result from modulating distributed CSTC loops rather than acting as a focal lesion or pacemaker at the stimulation site.

The lack of cognitive control improvement from midSTR-neuron opto-DBS also supports the clinical findings of Basu et al. and others [24,54,62]. Gray matter nuclei, such as the nucleus accumbens, have been investigated as targets for DBS therapies for OCD and addiction with mixed effects [3,63]. These targets can be advantageous over white matter tracts because they typically require lower stimulation amplitudes which extends battery life [11,63,64]. However, DBS of gray matter nuclei or stimulation through more ventral electrode contacts in the VCVS are associated with higher incidences of negative side effects like increased anxiety [43]. The striatum relies on a complex balance of direct/D1 and indirect/D2 MSNs and interneurons to gate cognitive control and other important decision-making components [65,66]. The midSTR-neuron opto-DBS used in the present study activates these different populations indiscriminately, which might lead to a “noisy” striatal output that is disadvantageous for cognitive control after a sustained period of activation. Further testing is needed to elucidate the process by which stimulation impairs cognitive control. These results also imply that to achieve the cognitive control benefit, electrical DBS must engage white matter fibers passing to and through the midSTR. Importantly, this does not rule out the possibility of a contributing role for local midSTR neurons in electrical DBS effects, which might explain our observation of attenuation in axonal opto-DBS effects over time.

A critical insight from this study was the temporal instability in the behavioral effects of opto-DBS. While the RT effects of electrical DBS are stable across days (Table S15), the cognitive enhancement from axonal opto-DBS faded within and across testing days. Electrophysiology results revealed network-level (polysynaptic) plasticity-like changes within and across days, but the mechanisms of these “fading” effects are still unclear.

Within a day, the attenuation of the RT improvement was linked to a reduction in local midSTR evoked responses. We hypothesize this decay reflects physiological constraints that are inherent to optical stimulation rather than a lack of construct validity. Although the Chronos opsin has fast kinetics [39,67–69], driving this circuit continuously at 130 Hz may exceed biological limits, for example, causing a depolarization block or depletion at synaptic terminals [70]. High-frequency electrical stimulation can also exceed these limits [71] but acts through different channels to alter ionic currents within neurons or axons, which might lead to different desensitization thresholds in response to the two stimulation modalities [41,72].

Across Set-Shift sessions, the reduction in axonal opto-DBS’s behavioral effect corresponded to attenuated mPFC responsivity. Opto-DBS’s effect on the early and late components of the mPFC response declined across days, and the reduction in the early component was accompanied by a decrease in the slope of the stimulation dose-response curve. In other words, the relative post versus pre opto-DBS difference decreased over the course of testing. This implies the mPFC became less excitable or desensitized by DBS, and hints at cortical plasticity mechanisms. Long-term potentiation is caused by high-frequency stimulation, like the 130 Hz used for DBS and can occur in corticostriatal pathways [73–75]. The early component of the mPFC ERP likely corresponds to glutamatergic signaling and is NMDA receptor-dependent [49,50,76]. This further supports the idea that the “fading” effect of axonal opto-DBS on cognitive control is driven by a plasticity-like process in excitatory mPFC neurons rather than at the terminals in midSTR. Future studies using opto-DBS may need to adopt intermittent stimulation paradigms (e.g. trial or block-based, [28]) rather than continuous high-frequency stimulation to mitigate these adaptations.

We acknowledge several study limitations. Anatomically, rats lack a defined internal capsule at the anterior-posterior level of the midSTR DBS target, meaning mPFC axons are intermixed with striatal neurons [10]. While our opto-DBS approach allowed us to functionally separate these components, the homology to the human internal capsule is imperfect. One solution is to test more posterior stimulation targets, where axons do come together to form a true capsule [77]. On the other hand, that more posterior target compresses fibers together and may not be able to replicate the human cognitive enhancement or lead to off-target effects.

Additionally, our electrophysiological investigation of the “fading” effect used a small sample size; however, the robustness of the ERP changes still provides compelling preliminary evidence for the neurophysiological mechanisms involved. As discussed above, high-frequency optogenetic stimulation may not be biologically capable of fully replicating electrical stimulation, but remains a powerful tool for investigating the effects of DBS delivery to specific neural circuits [32,72,78]. Future research should clarify if the temporal instability of opto-DBS’s effects is due to specific limitations of optogenetic stimulation or if it has mechanistic meaning for how DBS improves cognitive control, particularly for white matter targets [41].

Finally, while we primarily targeted the mPFC-midSTR circuit based on prior work by our group and colleagues [8,27,10,29,9] other axons passing through the VCVS might contribute to DBS’s effects. Projections from the orbitofrontal cortex, an area that is dysfunctional in OCD patients, frequently overlap with those from the mPFC, and both are likely to be engaged by midSTR electrical stimulation [10,46,48,76,79]. Interestingly, electrical DBS delivered to this pathway produced changes in LFP power in the orbitofrontal cortex that developed during the 30 min stimulation period [76], similar to the attenuation in cognitive effects and ERP magnitude observed here. That study did not observe any power changes in the mPFC; however, this may be due to the DBS location which was slightly posterior to our midSTR target. Further, our midSTR target also contains axons that do not terminate in the striatum but instead continue on to downstream structures like the thalamus [10]. These axons can still express the Chronos opsin but will likely be differently engaged by DBS compared to those with terminating in striatum [41]. Like the tractography-guided approaches in clinical settings [54,80], follow-up studies could deliver electrical or optogenetic DBS to these targets and evaluate their effects on cognitive control.

In conclusion, this study provides behavioral and physiological evidence that the cognitive control benefits of VCVS DBS are driven primarily through engagement of prefrontal axons in the internal capsule, not local striatal neurons. This implies that therapeutic success depends on engaging specific brain networks, not just activating (or inactivating) a general region of gray matter. Clarifying the origins of time-dependent changes in opto-DBS’s efficacy and how this model compares to electrical stimulation in rodents and humans will be important to solidify the translational validity of this and other preclinical DBS paradigms, ensuring continued clinical impact.

---

**Supplemental Figure 1:**
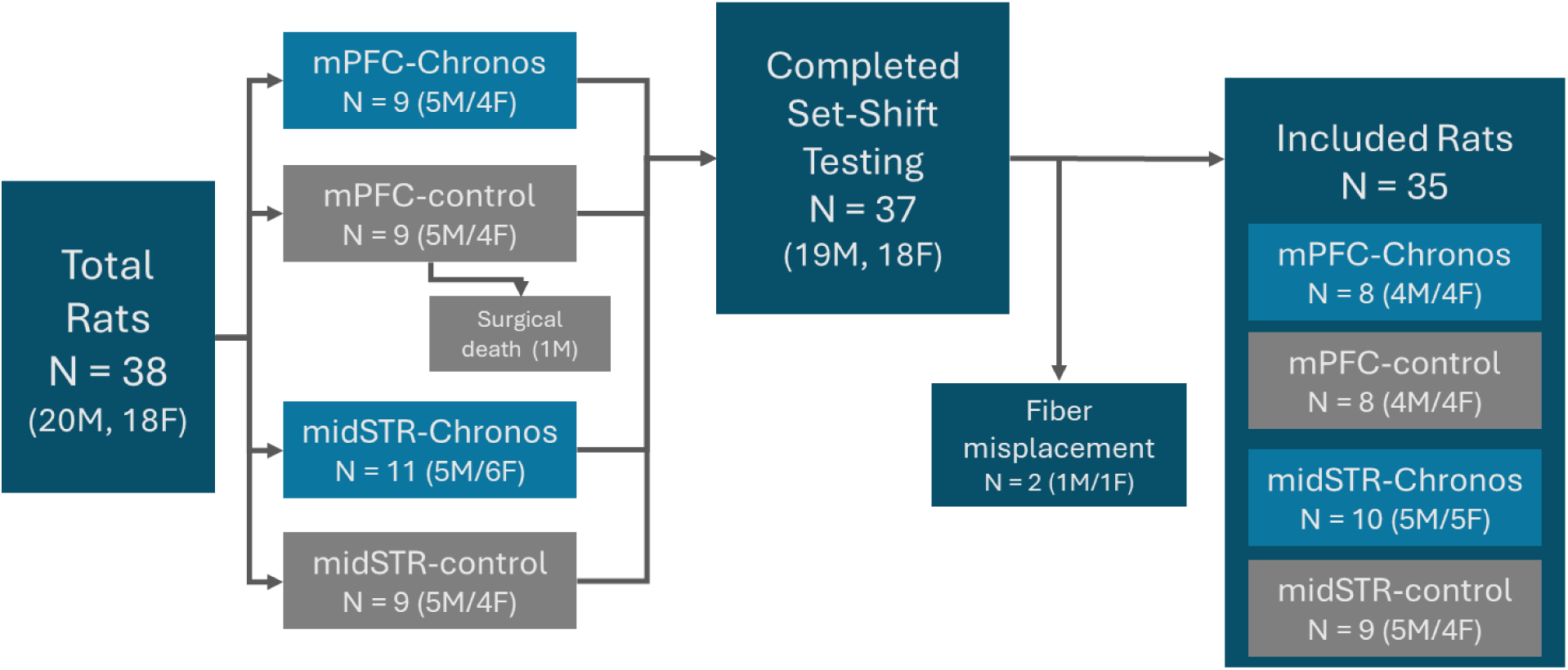
Consort Diagram

**Supplemental Figure 2:**
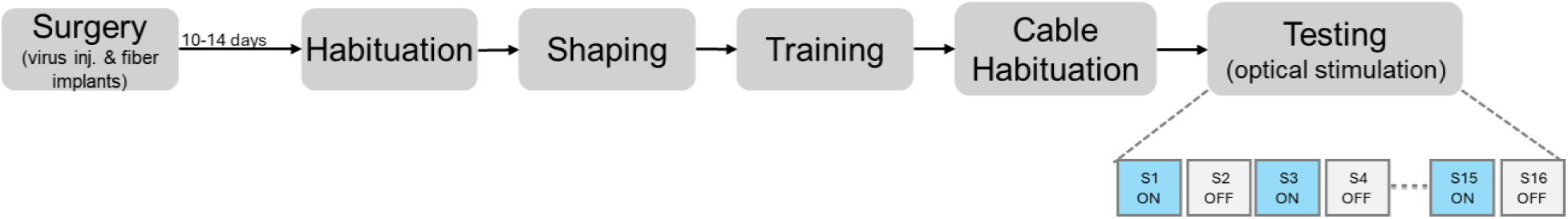
Set-Shift Experiment Schedule

**Supplemental figure 3:**
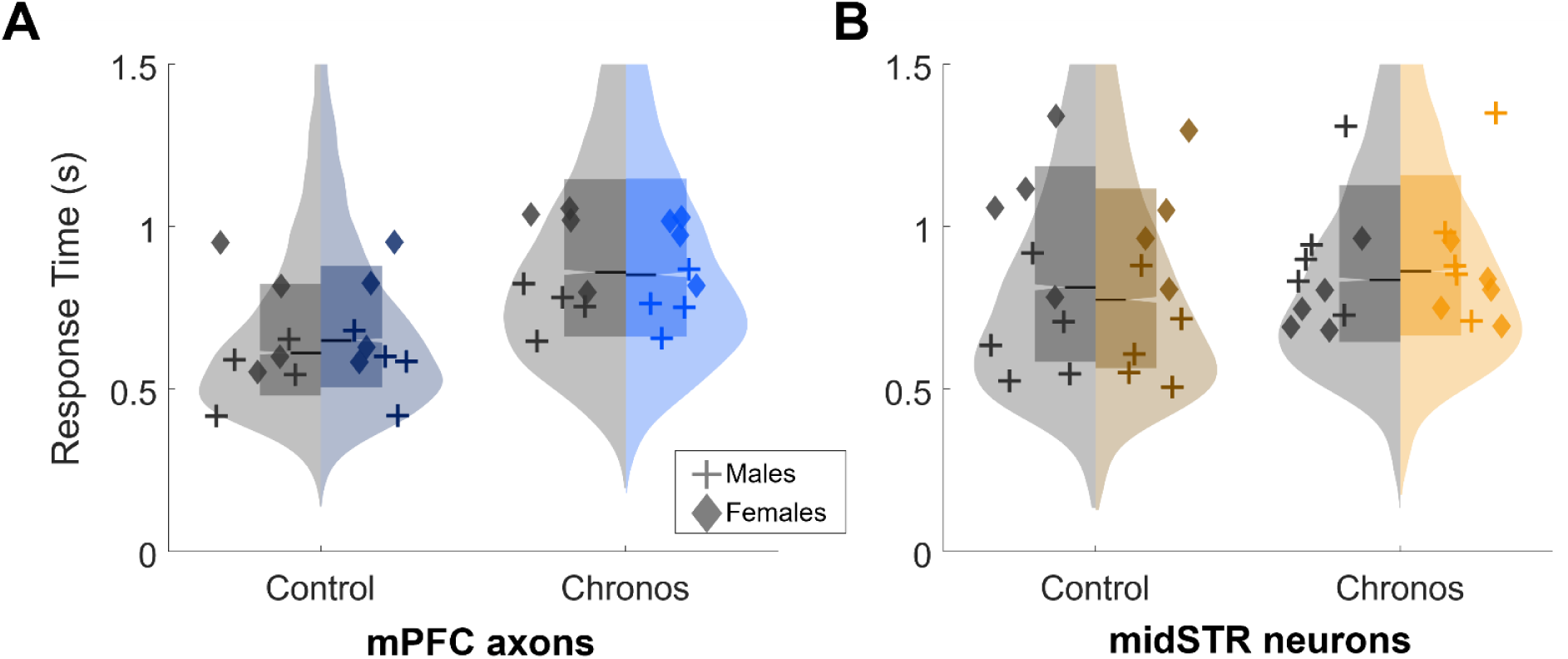
Opto-DBS ON and OFF raw response times

**Supplemental Figure 4:**
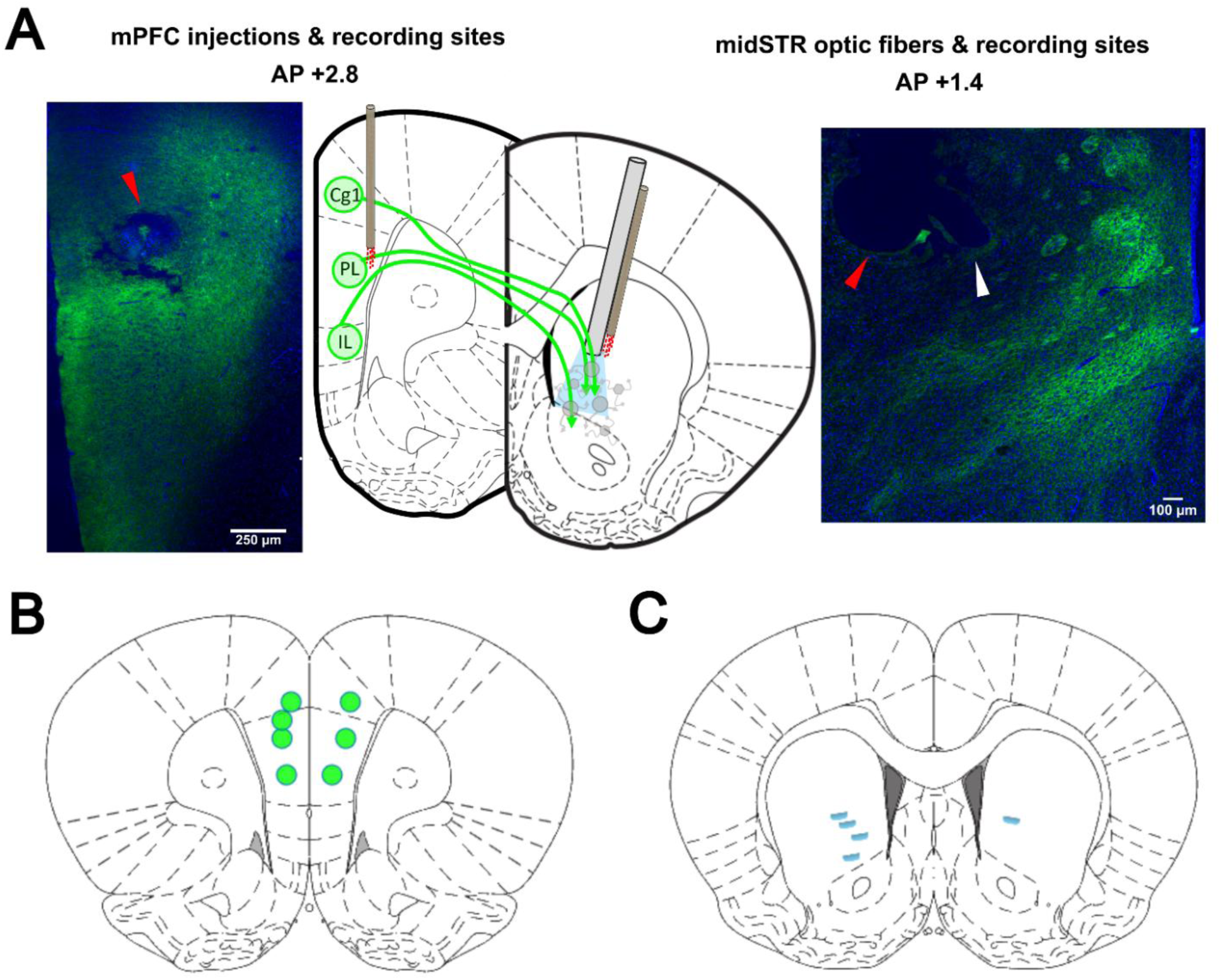
Evoked response potential experiment fiber and electrode locations **Setup of evoked-response potential experiments and histology results** **A) Left:** Red arrow points to example lesion site from a microwire used to record local field potentials in the mPFC, surrounded by Chronos-GFP+ neurons. **Middle:** Location of injection sites in mPFC subregions (Cg1, PL, and IL as in Paxinos & Watson 2007, or A24b/A32b, A24a/A32a, and A25, respectively, as in Paxinos & Watson 2013). Recording wires were targeted to PL/A24a/A32a. Optic fibers were implanted in midSTR, similar to Figure 2A, but here alongside recording microwires. **Right:** Location of fiber and recording site in the midSTR stimulation target positioned above Chronos-GFP+ axons and axon terminals. White arrow indicates fiber tip. Red arrow points to site of recording wire. **B)** Locations of 7 mPFC cortical recording sites (4 left hemisphere, 3 right) used for the ERP analysis. Atlas slice represents average AP coordinate (+2.87, Paxinos & Watson 2013). 5 sites were in PL (A24a/A32a), and 2 sites were in Cg1 (A24b/A32b). **C)** Location of 5 midSTR optic fibers and recording sites (4 left, 1 right) used for the ERP analysis. Atlas slice represents average AP of +1.59.

## Notes

### Competing Interest Statement

The authors have declared no competing interest.

### Funding

- National Institute of Neurological Disorders and Stroke: 1R01NS120851-01A1

